# Maize plant infection by *Ustilago maydis* is regulated by the Fungal Sulfur Metabolism

**DOI:** 10.1101/2025.03.13.642974

**Authors:** Emilio Espinoza-Simón, Nayeli Torres-Ramírez, Margarita Juárez-Montiel, Emmanuel Ríos-Castro, Damián Ramírez-Robles, Rosario Ortíz-Hernández, Olga Echeverría-Martínez, Lourdes Villa-Tanaca, Francisco Torres-Quiroz

## Abstract

The sulfur metabolism is tightly regulated in cells. Cysteine, at physiological concentrations, plays a crucial role in protein assembly, as well as in coenzyme and metabolic intermediate synthesis. Additionally, cysteine is biotransformed into H_2_S, a gasotransmitter with several roles on cells, ranging from regulating mitochondrial metabolism to producing metabolic intermediates and mediating post-translational modification of proteins. While H_2_S has been shown to participate in the infection processes of animal pathogenic fungi, its role in phytopathogenic fungi remains unexplored. Here, we describe the conditions required to induce endogenous production of H_2_S in plant pathogenic fungi *Ustilago maydis*. Under these conditions, we observed an increased infection rate and more pronounced symptoms in maize plants. A label-free proteomic assay to examine adaptations of *U. maydis* under H_2_S-producing conditions shown an increased expression of extracellular enzymes required for virulence and mitochondrial proteins related to cellular respiration, ATP synthesis, and fatty acid degradation, along with enhanced expression of proteins involved in proteasomal degradation. Conversely, we found reduced expression of proteins associated with antioxidant responses, glycolysis, and the pentose phosphate pathway. These mitochondrial protein alterations correlated with increased mitochondrial biogenesis, ultrastructural changes, inhibition of the cytochrome respiratory pathway, and elevated activity of an alternative oxidase. Additionally, H₂O₂ production increased, while the enzymatic capacity for its detoxification decreased. Impaired lipid accumulation and altered intracellular distribution were also observed. Thus, in *U. maydis*, the modulation of cysteine metabolism regulates mitochondrial function, protein expression, lipid metabolism and infectious processes.

**Author Summary:** *Ustilago maydis is a basidiomycete fungus that infects corn plants, inducing tumor formation. While in some countries this infection causes significant losses in maize crops, in Mexico, U. maydis, known as “huitlacoche,” is celebrated as a culinary delicacy with important nutritional value. Additionally, it serves as an interesting model for studying infection by dimorphic phytopathogenic fungi*.

*Animal models of fungal pathogenesis show that hydrogen sulfide (H_2_S), whether exogenously provided or induced by supplementation with its precursor cysteine, plays a role in infection processes in both the pathogen and the host cell*.

*In this study, we explored the role of cysteine in the morphology, metabolism, and pathogenicity of U. maydis. Our findings indicate that cysteine treatment triggers an overproduction of H_2_S, alters mitochondrial morphology and nitrogen metabolism, and disrupts the oxidative balance in U. maydis. Furthermore, fungi to cysteine enhance tumor formation and anthocyanins accumulation in Zea mays plants. These findings suggest that H_2_S may play a key role in the infection efficiency of phytopathogenic fungi*.

## Introduction

Regulation of sulfur metabolism is essential in cells. In yeast, sulfur is available in several chemical forms: sulfate, sulfites, thiosulfate, elemental sulfur, and organic species such as methionine, glutathione, homocysteine, and cysteine. Two metabolic pathways, sulfate assimilation and the transsulfuration pathway, regulate sulfur biochemical reactions. In the sulfate assimilation pathway, yeast uptake inorganic sulfides (sulfite, sulfate, or thiosulfate) and produce homocysteine, S-adenosyl-methionine, and methionine. In the transsulfuration pathway, yeast uses organic sulfur molecules, like glutathione, cysteine, and methionine [1]. Cysteine plays a key role in the synthesis of glutathione and coenzymes, the remethylation pathway, and protein synthesis. At concentrations higher than 5 mM, cysteine is highly toxic in yeast [2] due to its highly reactive thiol group, and vacuole plays a main role in compartmentalizing intracellular amino acids distribution. High cytoplasmic cysteine levels impair mitochondrial respiration, limiting iron availability [3].

Cysteine levels are kept low inside cells and are compartmentalized in vacuoles [3]. In the transsulfuration pathway, cysteine is converted to H_2_S by several enzymes, such as Cys4 (cystathionine beta synthase) and Cys3 (cystathionine gamma-lyase). Regulation of intracellular cysteine levels varies among yeast species. *Candida albicans* synthesizes sulfite from cysteine overload using several enzymes like cysteine dioxygenase, and then this is effluxed out of the cells using a sulfite efflux pump [4]. *Saccharomyces cerevisiae* instead lacks cysteine dioxygenase and metabolizes cysteine to cystine and other non-reactive thiols, like cystathionine, glutathione, and S-Adenosyl-L-homocysteine [5]. Furthermore, *S. cerevisiae* grown in rich cysteine media increases H_2_S production [6]. In *Ustilago maydis*, several hypothetical enzymes related to H_2_S metabolism have been described, such as cysteine synthase, cystathionine gamma synthase, sulfite reductase, and sulfide quinone oxidoreductase [7].

Both sulfur metabolism pathways produce H_2_S, a gasotransmitter with several roles in cells. This molecule plays a hormetic role in mitochondrial bioenergetics, stimulating ATP production at low concentrations while inhibiting the cytochrome pathway at high concentrations and decreasing cell respiration [8]. Also, H_2_S interacts with metalloproteins to regulate signal transduction, cellular metabolism, antioxidant response, and detoxification [9],[10]. Additionally, H_2_S induces redox post-translational modifications in protein cysteine residues, which could regulate protein structure and activity [11]. Among these modifications, S-persulfidation and S-sulfenylation regulate protein folding [12], signal transduction [13], and protein oligomerization [14].

H_2_S plays several roles among cells, from energy production and antibiotic resistance in bacteria [15], to photosynthesis and ripening in plants [16], as well as bone healing, neurotransmission, and insulin signaling in mammals [8]. In *Escherichia coli* and *Acinetobacter baumannii*, endogenous H_2_S increases respiratory flux, enhancing energy production and respiration [17], while in *Mycobacterium tuberculosis*, it enhances survival during mouse infection [18]. In infected cells, *Helicobacter pylori* induces host cystathionine gamma lyase (CSE) expression, thereby increasing cystathionine production and enhancing bacterial survival inside cells. Additionally, bacterial strains lacking CSE are easily cleared by infected macrophages [19]. Antibiotic resistance is linked to sulfur metabolism in bacteria, as demonstrated in *Pseudomonas aeruginosa*, *Staphylococcus aureus*, and *Bacillus anthracis*. It was shown that the inactivation of enzymes such as cystathionine beta-synthase, cystathionine gamma-lyase, or mercaptopyruvate sulfurtransferase makes these bacteria more susceptible [15]. Therefore, in a *ΔcstR Staphylococcus aureus* strain that cannot regulate persulfide levels, a decrease in virulence gene expression is observed, suggesting a role for persulfide levels in bacterial pathogenicity [20]

In *S. cerevisiae*, H_2_S has a dual role. At low concentrations, it enhances antioxidant activity upon heavy metal exposure [21], regulates glucose fermentation and oxygen consumption [22], growth [23], extends chronological lifespan [24],[25] and increases oxidative stress resistance [26]. However, at high concentrations, H_2_S inhibits mitochondrial oxygen consumption [27] and yeast growth [23]. Also, H_2_S regulates the ultradian cycle and population synchrony [28].

In *Aspergillus niger* and *Penicillium italicum*, H_2_S exposure impairs spore germination and mycelial growth [29]. Also, exposure to H_2_S inhibits the growth of the aforementioned fungi during fruit harvest [30]. Sueiro-Olivares showed that a low-S-persulfidation *Aspergillus fumigatus* strain has reduced virulence and a low fitness against host cell immune response. Also, a relationship between host cells and *A. fumigatus* S-persulfidation is closely related [31]. Besides, in *Candida albicans*, the H_2_S-producing enzyme cystathionine beta-synthase (CBS) plays a role in antioxidant response and pathogenicity. Also, pharmacological CBS inhibition decreases virulence in a murine candidiasis model [32].

*Ustilago maydis* is a dimorphic pathogenic basidiomycete that infests corn, causing tumors in aerial parts of the plant. This fungus exhibits a biphasic life cycle. Following spore germination in the maize plant epidermis, it initially grows as haploid saprophytic yeast, undergoing asexual reproduction [33]. However, upon mating, two haploid cells fuse, forming a pathogenic dikaryon that secretes lytic enzymes degrading the cell wall. Subsequently, the fungus propagates within vegetal cells. Tumor formation typically initiates five days after infection, marking the onset of the sporogenesis stage and producing teliospores [34]. *U. maydis* is considered a pathogenic fungus that causes losses in maize crops across the world [35]; however, in Mexico this fungus, called huitlacoche (from nahuatl: black charcoal) is highly rated in gastronomy. Furthermore, huitlacoche is an important source of vitamins and beneficial peptides [36].

The pathogenicity of *U. maydis* relies on several factors, such as ustilagic acids, mannosylerythritol lipids, ferrochromes, and polyketides, among others [37]. Additionally, trehalose, a non-reducing disaccharide and compatible solute, plays a crucial role in fungus stress resistance and maize virulence [38]. Moreover, *U. maydis* strains exhibiting enhanced oxidative stress resistance have shown reduced virulence [39]. Effector genes such as Pep1, Rsp3, See1, Tin2, and Lep1 also play key roles during infection [37]. A secretome analysis has shown that most of the 40 core effectors regulate pathogenicity. These effectors are related to nutrition, adhesion, detoxification, and shielding [40],[41]. Therefore, secreted enzymes like endo-xylanases [42] and proteases [43] are critical for facilitating infection by breaking down plant barriers and enabling fungal colonization.

Here, the role of cysteine in H_2_S production and its effects on *U. maydis* physiology were analyzed. Following cysteine incubation, there was an observed increase in H_2_S production, mitochondrial biogenesis, and basal respiration. Notably, cysteine preincubation significantly enhanced the pathogenicity of *U. maydis* on *Zea mays* plants, leading to more severe infection symptoms, such as tumor formation and anthocyanins accumulation. Label-free quantitative mass spectrometry revealed differential expression in over 400 proteins, including virulence-associated extracellular enzymes, mitochondrial and proteasome proteins, and enzymes involved in secondary metabolites, antioxidant response, lipid, amino acid, and glucose metabolism. In line with this, fluorescence and electron microscopy revealed an increase in the number of mitochondria as well as alterations in morphology. Additionally, cysteine stimulates cyanide-resistant respiration and alternative oxidase activity. Moreover, cysteine triggered a redistribution of lipid droplets. Therefore, we found that cysteine induces an increase in ROS production and a decrease in H_2_O_2_ elimination. Also, we showed that cysteine alters S-sulfenylation and S-persulfidation of proteins, suggesting a role for H_2_S-induced PTMs on *U. maydis* physiology. Thus, H_2_S plays an important role in *U. maydis* pathogenicity through mitochondrial biogenesis and metabolic rewiring.

## Materials and methods

### 2.1 Culture of Ustilago maydis strains

*U. maydis* FB-2 (haploid non-infective, kindly provided by Oscar Flores [44]) and *U. maydis* SG200 (haploid infective) [45] were used. The SG200 strain of *Ustilago maydis* has an active b locus, where the bW and bE homeodomains interact to form a functional heterodimer. This homeodomain complex activates the signaling cascade required for the filamentous pathogenic state, enabling host colonization and tumor formation in *Zea mays*. Both strains were grown in solid YPD media (1% yeast extract, 2% peptone and 2% dextrose) and were incubated routinely at 30°C. To know the optimal conditions to induce H_2_S endogen production, yeast were cultured in **YPD**: 1% yeast extract, 2% dextrose, 2% gelatine peptone, **YPLac**: 1% yeast extract, 2% lactate, 1% gelatin peptone, 2% (NH_4_)_2_SO_4_, 0.1% KH_2_PO_4_; pH 5.5, synthetic fermentative medium or **SD**: 0.17% yeast nitrogen base, 2% glucose, 0.5% (NH_4_)_2_SO_4_, 0.1% KH_2_PO_4_, Drop Out 1X (in g/L: adenine 0.2, arginine 0.2, asparagine 1.0, glutamic acid 1.0, leucine 0.6, lysine 0.2, methionine 0.2, phenylalanine 0.5, serine 3.75, threonine 2.0, tryptophan 0.4, tyrosine 0.3, valine 1.5, uracile 2.0 and histidine 2.0), or synthetic respiratory medium or **SLac**: synthetic complete medium: 0.17% yeast nitrogen base, 2% lactate, 0.5% (NH_4_)_2_SO_4_, 0.1% KH_2_PO_4_, Drop Out 1X, pH 5.5. The previously prepared media were prepared with or without 2 mM **Cysteine.**

### 2.2 H_2_S production

H_2_S production was evaluated using a lead acetate strip assay [46]. Briefly, *U. maydis* FB2 or SG200 were cultivated overnight (ON) at 30°C in YPD with shaking (250 rpm). Next, cells were grown in SD, SD plus cysteine (SDcys), SLac, or SLac plus cysteine (SLcys) at 0.2 initial O.D._600_. Then, tubes were grown overnight at 30°C in a rotating carousel at 50 rpm (Cel-Gro Tissue Culture Rotator, Thermo Scientific). H_2_S reacted with acetate lead stripes, forming a dark brown to black precipitate. The length and darkness of the stripe were proportional to the H_2_S amount produced by yeast. H_2_S rate production was measured with a methylene blue assay [47]. *U. maydis* cells were cultivated overnight at 30°C in YPD. Then, cells were grown in SD, SDcys, SLac or SLcys at 0.2 initial O.D._600_ in a 96-well plate with methylene blue (1 mg/mL in citrate buffer). Growth (600 nm) and H_2_S production (663 nm) over 6h at 30°C were measured in an Infinite 200 (TECAN, Life Sciences, Mannedorf, Switzerland).

### 2.3 Cell viability test

Once we have cultured *U. maydis* (both strains) under previously mentioned conditions, cells were harvested, washed, and resuspended in sterile H_2_O to an O.D._600_= 1 and serial dilutions were dropped in YPD agar. Plates were incubated at 30°C and observed at 24h and 48 h [48].

### 2.3 Mitochondrial staining

*U. maydis* FB2 were cultivated as mentioned early. Next, yeast were grown in SD, SDcys, SLac or SLcys ON at 30°C. Then, cells were incubated with 25 nM Mitotracker Green (Thermo Fisher Scientific, Waltham, MS, USA) for 30 min at 30°C. Stained yeast were mounted in agarose pads [49] and observed under a Nikon Eclipse E600 microscope (Nikon Corporation, Japan). Images were recorded with a Nikon DXM1200F digital camera (Nikon Corporation).

### 2.4 Lipid staining

Upon *U. maydis* FB2 being grown and treated as above mentioned, cells were washed with PBS 1X twice, and then cells were fixed with 2% paraformaldehyde in PBS for 15 minutes. Then, cells were washed with PBS 1x twice, and cells were permeabilized with Triton X-100 0.1% in PBS 1X for 5 minutes. Cells were washed and stained with 5 µM of BODIPY^TM^ 493/503 (Thermo Fischer Sci) in PBS for 30 minutes [50]. Cells were washed twice and mounted in agarose pads. Slides were observed under a Nikon Eclipse E600 microscope (Nikon Corporation, Japan). Images were recorded with a Nikon DXM1200F digital camera (Nikon Corporation).

### 2.5 Electron microscopy

*U. maydis* FB2 cells, treated as described above,were fixed in 4% glutaraldehyde in 0.2 M PIPES buffer (pH 6.8) containing 0.2 sorbitol, 2 mM MgCl_2_ and 2 mM CaCl_2_, overnight at 4°C. Next, samples were rinsed with water and post-fixed in 2% KMnO_4_ for 45 min at room temperature. Cells were contrasted with 1% uranyl acetate and dehydrated in graded ethanol series from 30% to 100%. After dehydration, samples were infiltrated and embedded with Durcupan™ ACM (Sigma-Aldrich). Ultrathin sections of 60 nm were cut using the ultramicrotome Leica Ultracut UCT, mounted on copper grids covered with formvar and contrasted with uranyl acetate and lead citrate. Sections were observed under a Jeol 1010 electron microscope at 80 kV. Digital images were captured with a Hamamatsu camera (Hamamatsu Photonics K. K., Japan).

### 2.6 Sample preparation for LC-MS analysis

*U. maydis* FB2 cells were cultured in SLac or SLcys as mentioned above. Then, cells were lysed in liquid nitrogen using glass beads, mortar and pestle. Cell debris was separated by centrifugation at 14000 rpm for 15 min at 4°C. Next, proteins were precipitated using CHCl_3_/MeOH. Then, proteins were dissolved in PBS 1X and protein was quantified by the Pierce BCA Assay Kit (Thermo Fisher Scientific, Waltham, MS, USA).

The supernatant proteins were precipitated with methanol-chloroform, and the resulting pellets were enzymatically digested using the iST Sample Preparation iST® kit (PreOmics, Munich, Germany) according to the protocol established by the manufacturer. The enzymatic digestion was performed for 2 h in a Thermoblock (Eppendorf, Hamburg, GER). The resulting peptides were dried using a Savant DNA120 SpeedVac Concentrator (Thermo Fisher Scientific, Waltham, MS, USA) and then resuspended with the “LC-load” reagent (PreOmics, Munich, Germany). Finally, peptides were stored at –20°C until LC-MS analysis.

### 2.7 Mass spectrometry-based quantitative proteomics

Proteomic analysis was performed in a UPLC Nano Acquity M-Class coupled with a QTOF Mass Spectrometer Synapt G2-S*i* (Waters Corporation; Milford, MA, USA), according to the modified method of Ríos-Castro [51]. Briefly, tryptic peptides were separated on an HSS T3 C18 column (Waters, Milford, MA); 75 μm × 150 mm, 100 Å pore size, 1.8 μm particle size; using a mobile phase A, 0.1% formic acid (FA) in H_2_O, and mobile phase B, 0.1% FA in acetonitrile (ACN) under the following gradient: 0 min 7% B, 121.49 min 40% B, 123.15 to 126.46 min 85% B, 129 to 130 min 7% B, at a flow of 400 nL·min^−1^ and 45°C in the column compartment. The spectra data were acquired using a data-independent acquisition (DIA) approach through High-Definition Multiplexed MS/MS (HDMS^E^) mode (Full-Scan DIA). Parameters on ionization source were set with the following values: 2.75 kV in the capillary emitter, 30 V in the sampling cone, 30 V in the source offset, 70°C for the source temperature, 0.5 bar for the nanoflow gas, and 150 L·h^−1^ for the purge gas flow. Low and high-energy chromatograms were acquired in positive mode in a range of *m/z* 50−2000 with a scan time of 500 ms. Precursor ions were fragmented in the transfer cell using a collision energy ramp from 19 to 55 eV. Synapt G2-S*i* was calibrated with [Glu1]-fibrinopeptide, [M+2H]^2+^ = 785.84261 at less than 1 ppm across all MS/MS measurements.

### 2.8 MS-Data Analysis

The MS and MS/MS spectra contained in the generated *.raw files were quantified by Progenesis QI for Proteomics software *v*4.2.1 [52], [53] (Nonlinear Dynamics, Ann Arbor, MI, USA) using a target decoy strategy against a *Ustilago maydis* *.fasta database (downloaded from UniProt, UP000000561, 6805 protein sequences) [54], [55]. The parameters used for database search were: trypsin as a cut enzyme and one missed cleavage allowed; carbamidomethyl (C) as a fixed modification and oxidation (M) as a variable modification; peptide and fragment tolerance were set to automatic; minimum fragment ion matches per peptide: 2, minimum fragment ion matches per protein: 5, minimum peptide matches per protein: 1, and false discovery rate ≤ 1%. Only peptides between ±20 ppm were used for identification and quantification; besides, all false positive identifications (Reversed proteins) were discarded for subsequent analysis. Proteins considered differentially expressed presented at least a ratio ±1 (expressed as a base 2 logarithm); these proteins had at least ±2-absolute fold change and a p-value ≤ 0.05. The ratio was calculated by dividing the average MS signal response of the three most intense tryptic peptides (Top3) [56] of each well-characterized protein in the SLcys condition by the Top3 of each protein in the SLac condition. The mass spectrometry proteomics data have been deposited to the ProteomeXchange Consortium via the PRIDE [57] partner repository with the dataset identifier PXD060479. Results were exported to a *.csv file, which contained all MS measurements (supplementary file), this file was analyzed in OmicScope [58] to determine and visualize potential DEPs

### 2.9 Oxygen consumption assay

*U. maydis* FB2 oxygen consumption was measured by high-resolution respirometry. Yeast cells were cultured as mentioned above. Then, cells were washed twice in distilled water and resuspended in a phosphate buffer (10 mM, pH 7.4). Oxygen consumption was determined using the Oroboros Oxygraphy-2k (Oroboros Instruments, Innsbruck, Austria) calibrated at 30°C. Basal respiration and, upon 1 mM glucose, 1 mM KCN, and 6 μM of n-Octylgalate (nOG) additions were measured.

### 2.10 S-persulfide and S-sulfenyl detection in U. maydis proteome

*U. maydis* FB2 proteome S-Sulfenylated was observed by SDS-PAGE of labeled protein with DAz-2:Cy5. Cells were cultured as mentioned above, and proteins were extracted with lysis buffer (50 mM HEPES pH 7.7, 1 mM EDTA, 0.1 mM neocuproine, 3% NONIDET P-40, 5% SDS, 3% Triton X-100, 1 mM PMSF, 1X protease inhibitor) adding 100 μM of DAz-2:Cy5. This solution was dropped into liquid nitrogen in a mortar with 425-600 µm glass beads. Once it was frozen, the solution was pulverized with a pestle. The resulting solution was transferred to a 1.6 mL amber microtube and incubated at 37°C for 90 min light protection. Then, proteins were separated from the beads by centrifugation (5 min, 3,100 rpm, 4°C) and the supernatant was transferred to another microtube, adding 4/4/1 portions (sample/methanol/chloroform) and centrifuged (15 min, 14,000 rpm at 4°C). Both aqueous and organic phases were eliminated, and proteins were washed twice with cold methanol and centrifugation (5 min, 14,000 rpm, 4°C). Finally, the pellet was resuspended in a sample buffer (50 mM HEPES pH 7.7 and 2% SDS). Samples were quantified by BCA assay, elucidated in a 12% SDS-PAGE and observed in a Typhoon FLA 9500 using an LPR filter for Cy5 (635 nm).

*U. maydis* proteome S-Persulfidated was observed by SDS-PAGE of labeled protein with NBF-Cl and DAz-2:Cy5. Cells were cultured and lysed as mentioned above, replacing DAz-2:Cy5 with 15 mM NBF-Cl. Once the solution was pulverized, it was incubated at 37°C for 60 min light-protected. Glass and cell components were separated from the proteins as mentioned before and resuspended in sample buffers (50 mM HEPES pH 7.7 and 2% SDS). Then, proteins were quantified with BCA assay, and a solution of 3 mg/mL of proteins was incubated with 100 µM DAz-2:Cy5 for 90 min. Methanol and chloroform were added to the same proportions mentioned above (4/4/1) centrifuged and washed twice with methanol. Proteins were resuspended in sample buffers, quantified by BCA assay, elucidated in a 12% SDS-PAGE and observed in a Typhoon FLA 9500 using an LPB filter for NBF (473 nm) and an LPR filter for Cy5 (635 nm).

### 2.11 H_2_O_2_ production

*U. maydis* FB2 were cultured as mentioned above. Upon incubation in SD, SD cys, SLac and SLcys, cells were centrifuged, and supernatant was collected to quantify H_2_O_2_ content. Cells were washed 3 times with distilled water and resuspended in PBS 1X. In a 96 well plate, supernatant or cells were added, and then 50 µL of Amplex Red (Thermo Fisher Scientific) mixture reactions were added. Yeast cells were incubated in an Infinite 200 (TECAN, Life Sciences, Mannedorf, Switzerland) for 3h at 30°C and fluorescence (530 exc and 590 emm) was recorded.

### 2.12 Catalase activity

To determine the role of H_2_S induced by cysteine on antioxidant enzymes, we measured catalase activity [59]. *U. maydis* FB2 cells were grown and treated as mentioned above. Cytoplasmic extracts were mixed with PBS and 1 mM H_2_O_2_ and incubated at 30°C for 5 minutes. Then, a work solution (Co(NO_3_)_2_ 5 mM in 1M NaHCO_3_) was added, vortexed for 5 sec, and incubated in darkness for 5 minutes. The absorbance at 440 nm was determined and is reported in U/mL*min.

### 2.13 Plant infection assay

Plant inoculation and symptom determination were performed to determine the role of cysteine-induced H_2_S in infectious processes. The method described by Juárez-Montiel, et al., (2015) was followed [60]. Maize seeds (cv. Cacahuazintle) were sown in pots with vermiculite, and after one week of germination, seedlings (about 45 per experiment) were inoculated with a syringe and needle into the stem with the solopathogenic *U. maydis* SG200 strain [7]. Plants were maintained in greenhouse conditions with a watering regime alternating water or nutrient solution h-e-mx, prepared according to the manufacturer’s instructions.

As previously described, the suspension of *U. maydis* SG200 sporidia was earlier treated with and without cysteine. Sporidia suspension was adjusted at O.D._600_= 1 in sterile water, and each of the 45 seedlings per condition was inoculated with 100 mL of suspension. Seedlings injected with water comprised the mock control [61]. After 12 days post-inoculation (dpi), symptoms were registered.

### 2.14 Statistical analysis

Data analysis was performed in GraphPad Prism version 8.02 (GraphPad software, Inc. La Jolla, CA, USA). Student T-test or one-way ANOVA (Tukey’s multiple comparison test) was used to analyze data. p<0.05 was defined as a significant difference.

## Results

### Non-fermentable synthetic medium supplemented with cysteine (SLcys) induces H_2_S production

Previous reports have shown that low nitrogen concentration [62] and high cysteine concentration [63] increase H_2_S production in *S. cerevisiae*. To investigate the effect of endogenous H_2_S production on the physiology and pathogenicity of *U. maydis*, we first assessed the conditions that induce the production of this gasotransmitter. To evaluate this, we cultured cells in different media: YPD, YPD-Cys, YPLac, YPLac-Cys, SD, SD-Cys, SLac and SLac-Cys. After 24 hours of culture, compared to rich media, the non-fermentable synthetic medium supplemented with 2 mM cysteine (SLcys) induced the highest H_2_S production (Fig 1A) without affecting cell viability (Supp Fig 1). Even at shorter times (6h), SLcys was the condition that enhanced H_2_S production (Fig 1B). These experiments were performed using the *U. maydis* strains FB2 and SG200, and both exhibited the same behavior under these conditions (Supp Fig 2).

**Figure 1.**
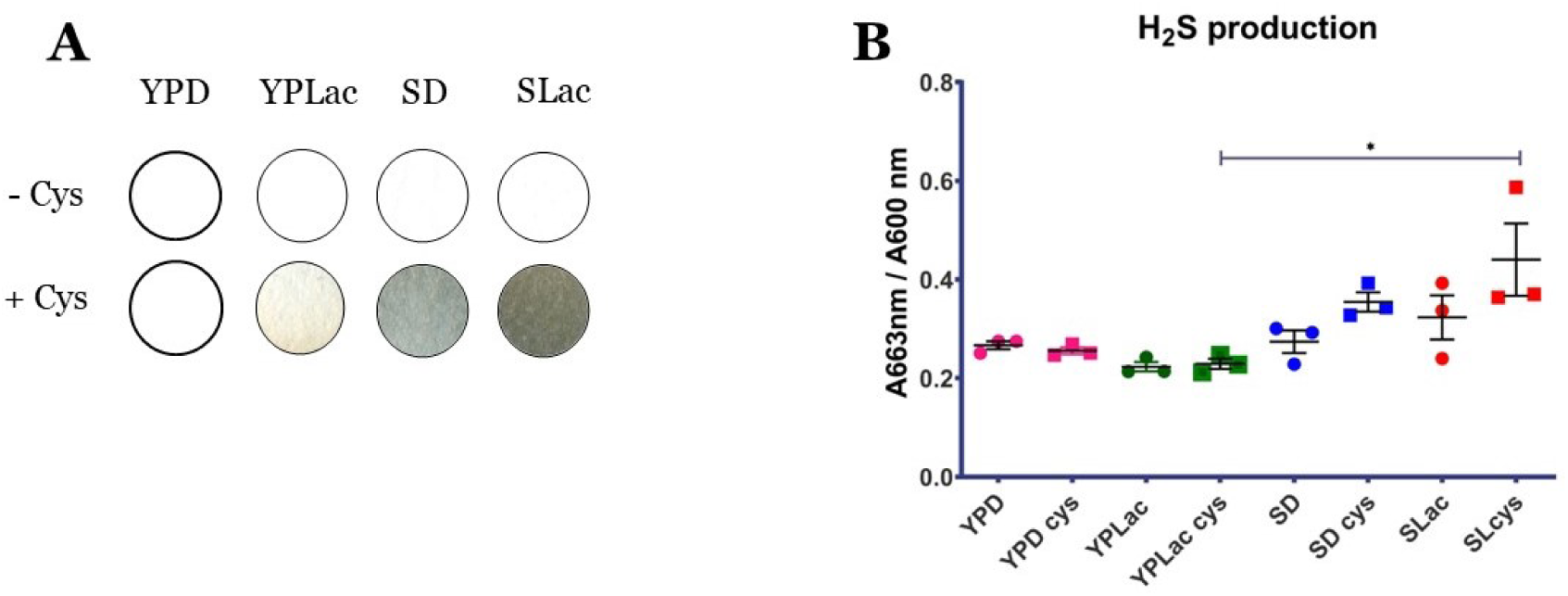
H_2_S production in *U. maydis* increases in respiratory media. *U. maydis* was cultured in YPD, YPD-Cys, YPLac, YPLac-Cys, SD, SD-Cys, SLac and SLac-Cys and H_2_S was detected with lead stripes after 24 hrs culture (Fig A) and methylene blue (Fig B) after 6 h incubation (* p<0.05).

**Figure 2.**
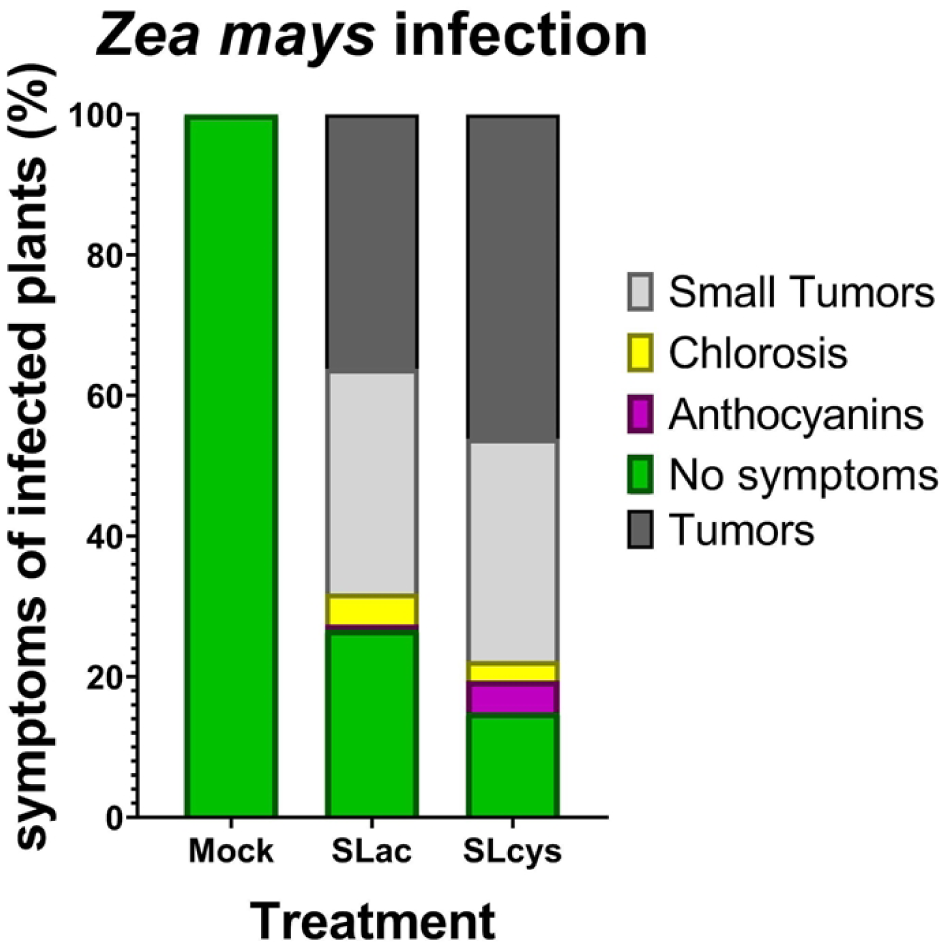
Infection of maize plants with *U. maydis* SG200 increased when yeast cells were supplemented with cysteine, with symptoms appearing 12 days post-inoculation.

In *S. cerevisiae*, incubation in fermentative media favors H_2_S production [22], however, in *U. maydis*, respiratory sources replicate the aforementioned phenomenon. This suggests that H_2_S production may be affected by the pathways involved in carbon utilization and energy production.

### Sulfur metabolism increases *Ustilago maydis* pathogenicity in *Zea mays*

Cysteine was observed to induce H_2_S overproduction, which in some pathogens is known to regulate virulence. Thus, we assessed the pathogenicity of the *Ustilago maydis* SG200 strain cultivated with cysteine in maize plants. Twelve days post-infection, an increase in plant developing tumors was observed (46.13% vs 36.17%, p<0.05), alongside an increase in the anthocyanins formation (4.53% vs 0.73%, p<0.05). (Fig 2 and Supp Fig 3). These findings highlight the dual role of cysteine in modulating pathogen behavior, underscoring its potential to influence the balance between virulence and plant survival in *U. maydis*.

**Figure 3.**
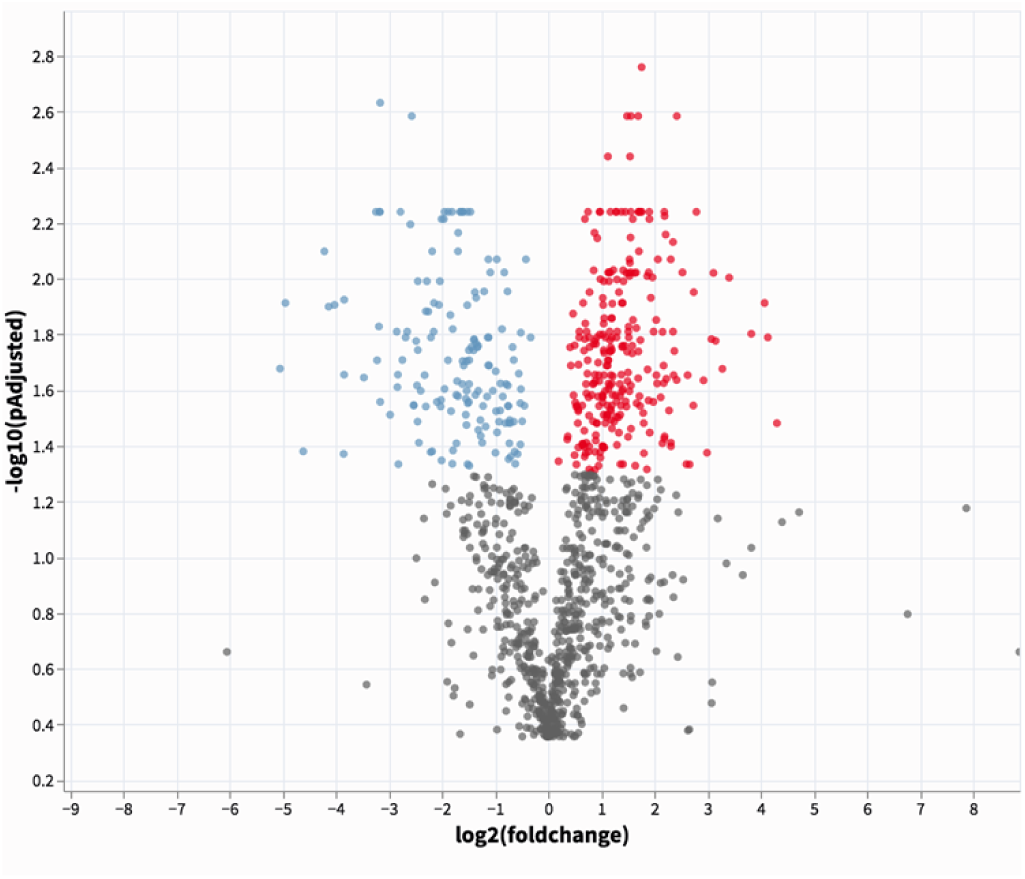
Cysteine induces differential protein expression in *U. maydis*. Upon cysteine exposure, cytoplasmic extracts were obtained, and the protein profile was characterized with “label-free” protein spectrometry. In this volcano plot, proteins upregulated are shown in red and proteins downregulated in blue (p<0.05). Graph obtained in Omicscope [58].

### Cysteine rewires the protein profile in *U. maydis*

Following the identification of alterations in H_2_S metabolism and *Ustilago* virulence after cysteine stimulation, we sought to investigate the modifications occurring at the proteomic level in *U. maydis*. A label-free proteomics experiment was conducted, comparing SLac and SLcys media. Quantitative analysis detected a total of 1,706 proteins, revealing distinct protein expression patterns between the two conditions (Fig. 3 and 4). These findings suggest significant alterations in metabolic mechanisms in cysteine-treated samples compared to untreated ones. Upon cysteine incubation, the expression of at least 400 proteins was dysregulated. Approximately 290 proteins were upregulated, with at least 190 showing a twofold increase in expression, while 160 proteins exhibited reduced expression, with at least 125 being downregulated twofold under the specified conditions (Supp Table 1 and 2).

**Figure 4.**
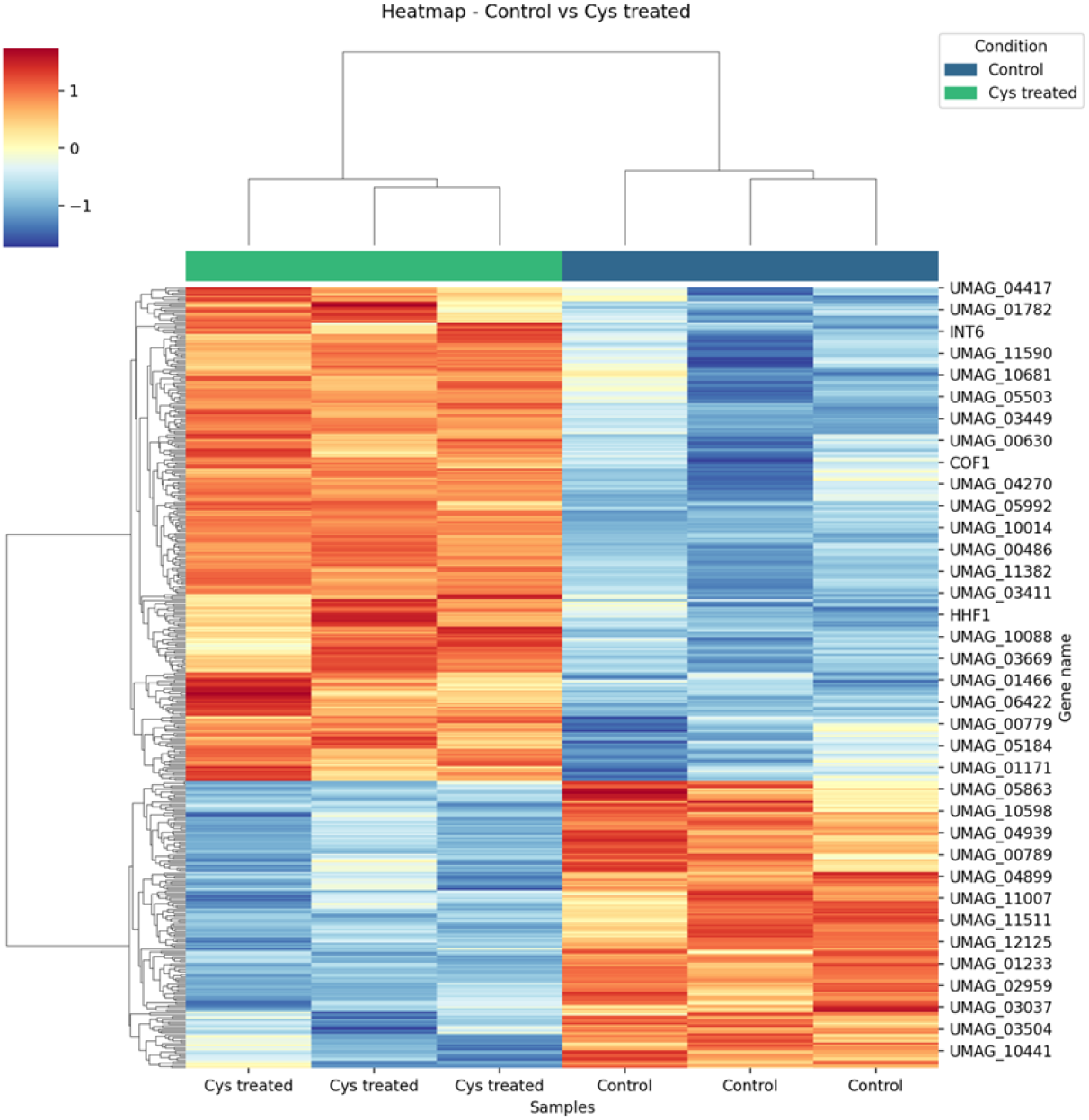
Cysteine induces differential protein expression in *U. maydis*. Upon cysteine exposure, cytoplasmic extracts were obtained, and the protein profile was characterized with “label-free” protein spectrometry. In this heatmap, red indicates high protein expression and blue low expression, with samples and proteins grouped by similarity. These results highlight specific molecular changes in response to cysteine treatment. Graph obtained in Omicscope [58].

Cysteine exposure triggered an increase in proteins related to mitochondrial processes, including respiratory electron transport, ATP synthesis, the mitochondrial respirasome, the TCA cycle, and fatty acid degradation. Additionally, nitrogen metabolism was significantly affected; proteins associated with amino acid biosynthesis, organonitrogen compound metabolism, and purine ribonucleotide metabolic processes were upregulated. Proteins linked to the proteasome complex also showed increased levels upon cysteine exposure. Conversely, proteins associated with sulfur compound metabolism, cellular responses to oxidative stress, the pentose phosphate pathway, glycolysis, the aldo/keto reductase family, and structural constituents of translational elongation were downregulated (Fig. 5).

**Figure 5.**
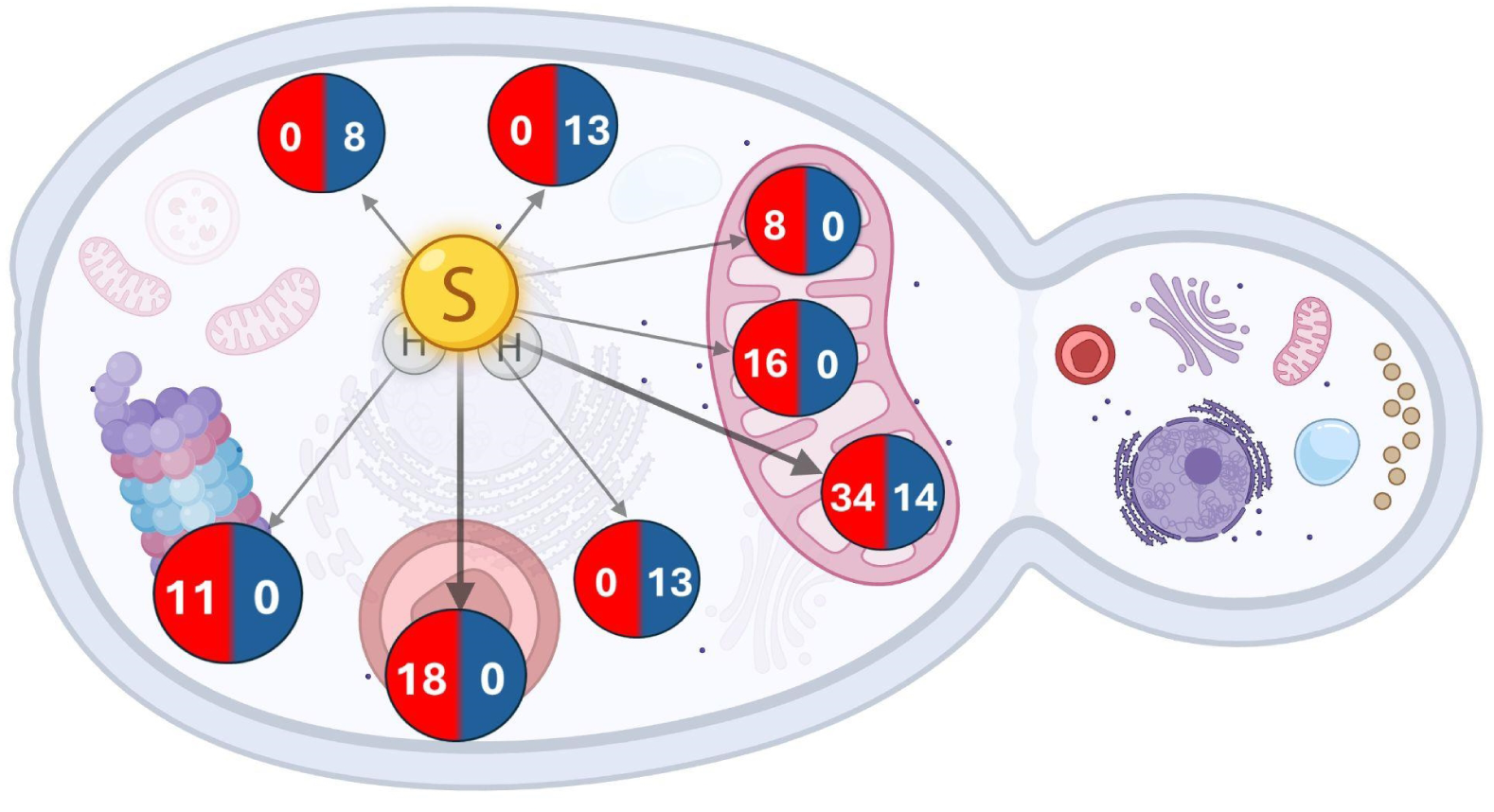
Proteome remodeling upon cysteine exposure in *U maydis*. Cysteine supplementation induces H₂S synthesis, leading to a reprogramming of the proteome. Mitochondrial, lysosomal, and proteasomal proteins are upregulated, while cytoplasmic proteins involved in carbon metabolism and protein synthesis are downregulated. Circles indicate proteins with increased (red) or decreased (blue) expression. Figure created with BioRender.com

Notably, cysteine incubation resulted in a threefold increase in endo-1,4-beta-xylanase levels and a striking ninefold increase in extracellular metalloproteinase levels, underscoring their potential roles in virulence modulation.

### Cysteine induces morphological alterations and an increase in mitochondrial biogenesis

The previously described results demonstrate that lactate, a mitochondrial energy supplier, combined with cysteine, induces increased H₂S production and significant protein profile rewiring, characterized by higher mitochondrial protein expression. Thus, to determine the effect of the aforementioned conditions on mitochondrial function, mitochondrial biomass, and morphology, they were evaluated using Mitotracker Green. In YPD or YPD-Cys media, abundant small mitochondria were observed, with a slight increase noted upon cysteine addition to the culture medium (Fig 6A and 6B). In synthetic fermentative media, mitochondrial biomass was low regardless of cysteine addition (Fig 6C and 6D). Growth in YPLac induces mitochondrial biogenesis, with a higher quantity of reticular-shaped mitochondria observed (Fig 6E). Interestingly, incubation in YPLcys induces a greater accumulation of mitochondria (Fig 6F), as well as the formation of pseudohyphae. Incubation in SLcys promotes higher mitochondrial biogenesis than SLac cultures, which apparently have an elongated shape (Fig 6H). These results indicate that the growth of *U. maydis* in nonfermentative media supplemented with cysteine affects mitochondrial morphology.

**Figure 6.**
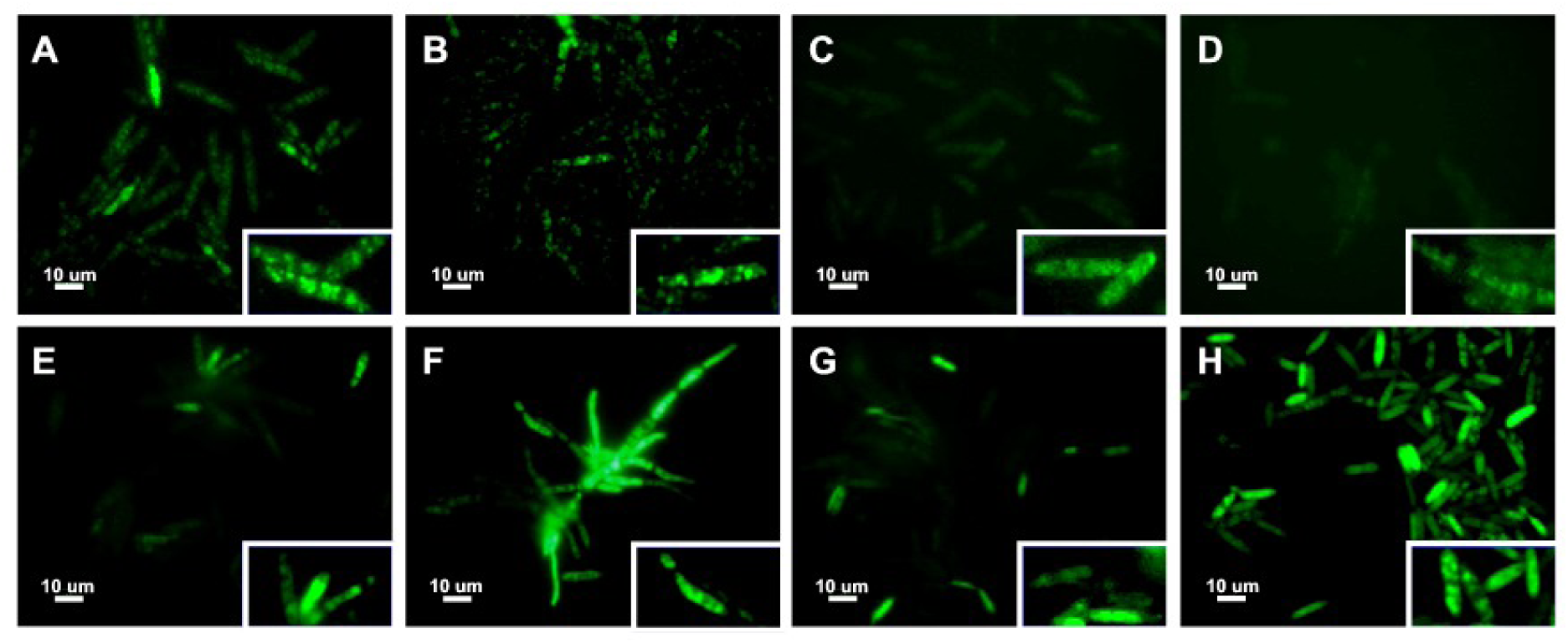
Cysteine induces mitochondrial biogenesis. *U. maydis* mitochondria were stained with mitotracker green after 24 hrs incubation in YPD (A), YPD-Cys (B), SD (C), SD-Cys (D), YPLac (E), YPLac-Cys (F), SLac (G) and SLcys (H). The insets show magnified regions where both fluorescence intensity and size are arbitrarily adjusted for visual clarity.

To characterize the morphological effects of the established culture conditions, yeast cells were studied using TEM. We observed that incubation with cysteine stimulates mitochondria formation, particularly in SLcys, where mitochondria appear more elongated (Fig 7H). Furthermore, cysteine reduces the formation and/or accumulation of lipid droplets. These findings demonstrate that incubation with cysteine, in addition to stimulating H_2_S production, has effects on cellular morphology and likely cellular metabolism.

**Figure 7.**
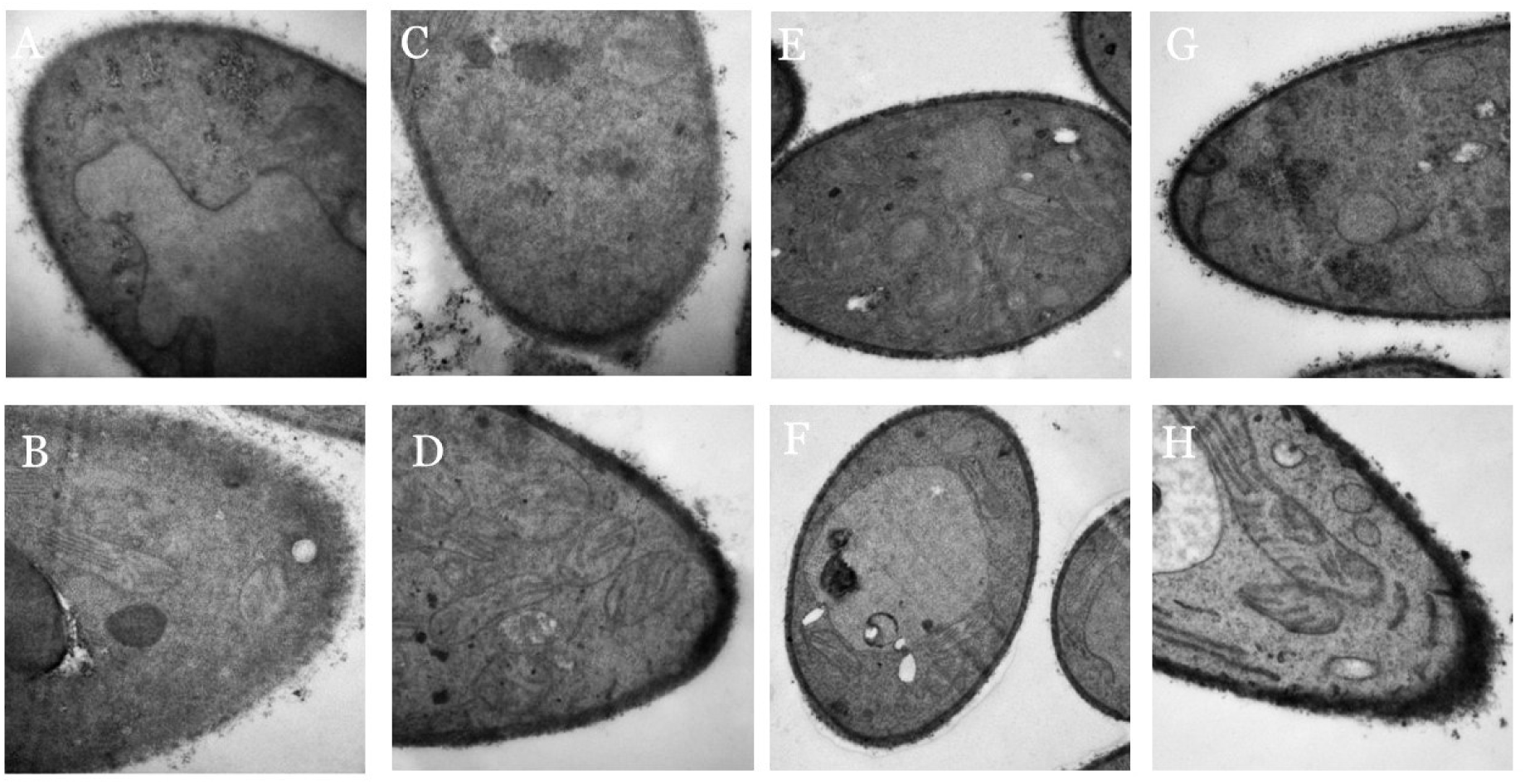
Cysteine induces mitochondrial ultrastructural changes *U. maydis* were analyzed with transmission electron microscopy after 24 hrs incubation in YPD (A), YPD+cys (B), YPLac (E), YPLac+cys (F), SD (C), SD+cys (D), SLac (G) and SLcys (H).

### Cysteine Decreases Cytochrome Respiratory Pathway Activity

Considering the evidence suggesting that cysteine significantly influences mitochondria, we characterize mitochondrial respiration under these conditions. As shown in Fig. 8, cysteine generally induces an increase in mitochondrial respiration, and interestingly, cysteine stimulates cyanide-resistant respiration, which suggests the presence of an alternative oxidase. This was confirmed by using a pharmacological inhibitor of this enzyme (Octyl gallate).

**Figure 8.**
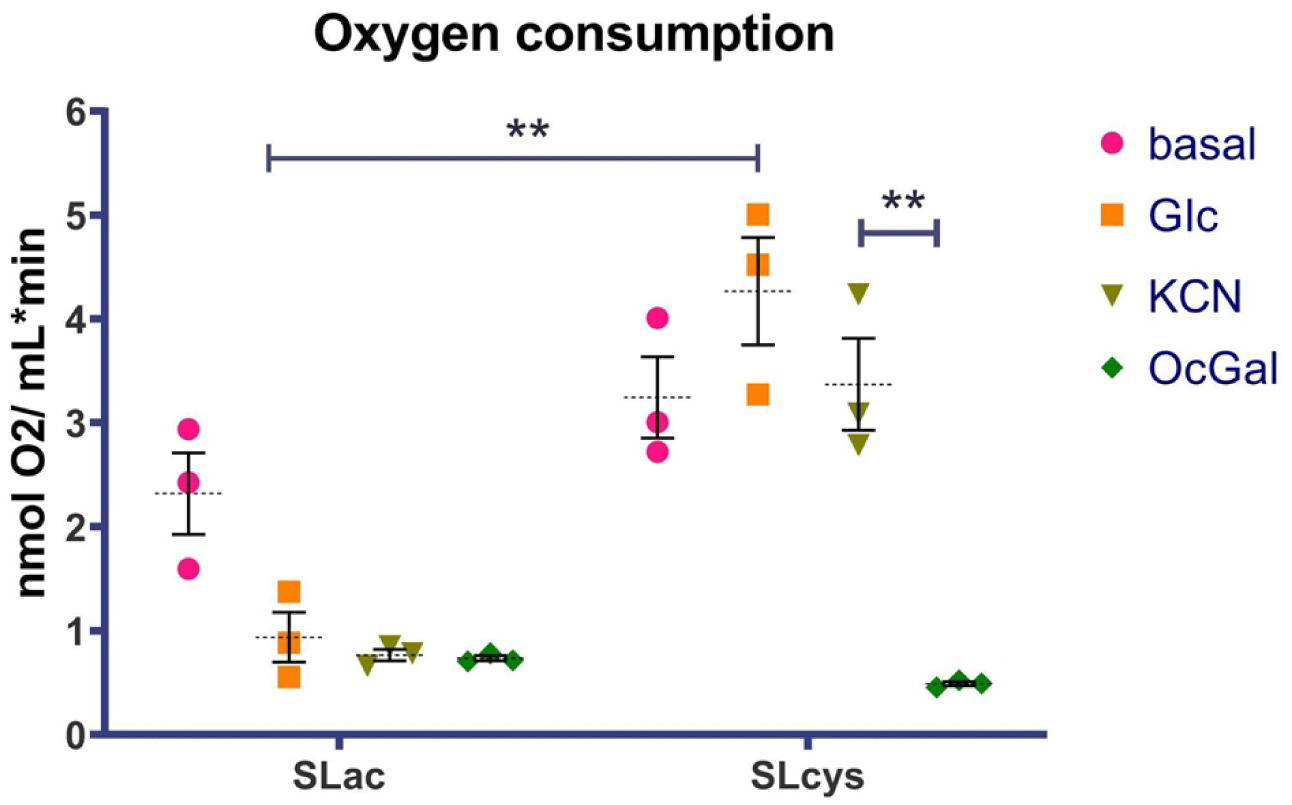
H_2_S impairs the cytochrome respiratory pathway. Oxygen consumption was measured in *U. maydis* under basal conditions of 1 mM glucose, 1 mM KCN and 6 μM *n*-Octylgalate (** p<0.005).

### H_2_S induced by Cysteine Impairs Antioxidant Activity in *U. maydis*

Our proteomic results showed that *U. maydis* grown in cysteine media expresses fewer antioxidant proteins. Thus, we measured H_2_O_2_ production in cells incubated under the aforementioned conditions. We observed that cysteine stimulates the accumulation of H_2_O_2_ in the medium after 24 hours of incubation (Fig. 9B) and also facilitates H_2_O_2_ production in yeast with fresh medium (Fig. 9A). Therefore, a decrease in H_2_O_2_ elimination in SLcys incubated cells was observed (Suppl Fig. 4). In summary, the addition of cysteine significantly alters mitochondrial metabolism by inhibiting the cytochrome pathway, enhancing alternative oxidase activity, promoting biogenesis, likely enabling mitochondrial fusion, and generating oxidative stress.

**Figure 9.**
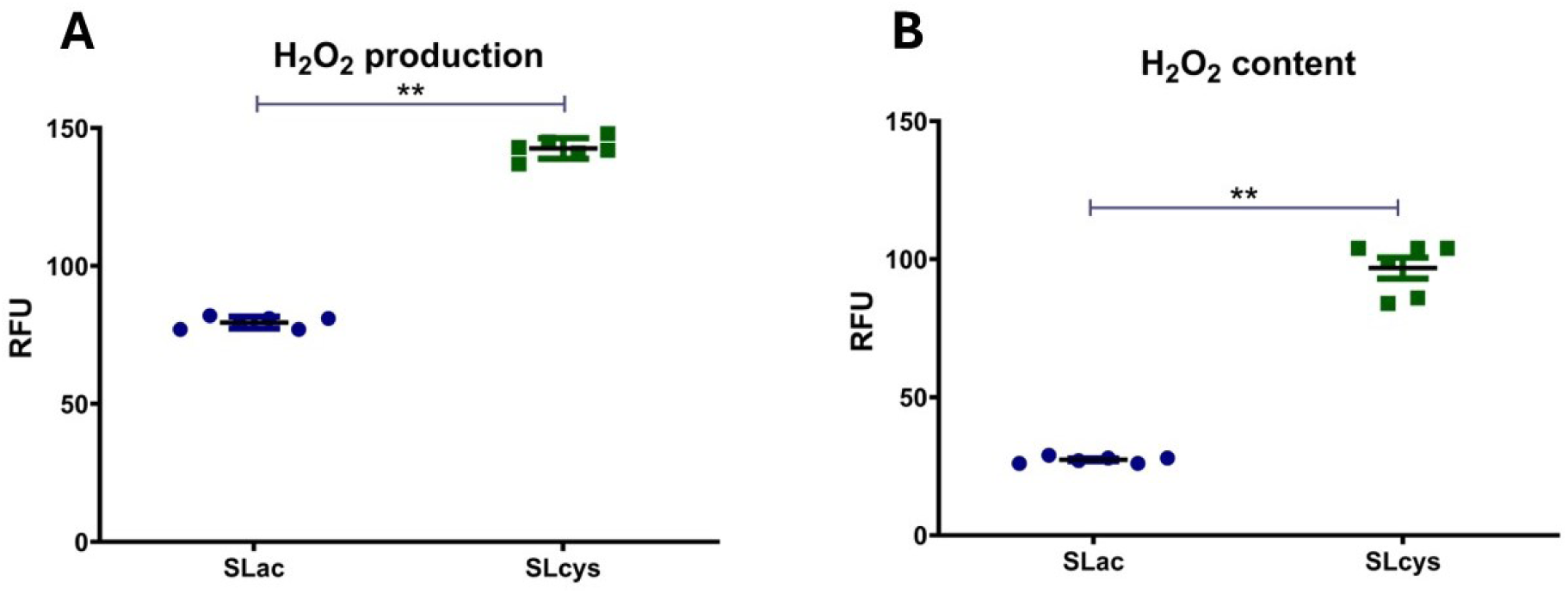
Cysteine preincubation raises ROS production. *U. maydis* grown with 2 mM Cysteine shows an increase in H_2_O_2_ accumulation in culture media (B). Also, upon changing the culture media, Cysteine-grown yeast showed higher H_2_O_2_ production (A) (p< 0.005).

### Cysteine Alters Lipid Distribution in *Ustilago maydis*

Since mass spectrometry results showed alterations in the production of proteins related to lipid metabolism, we decided to evaluate the lipid content and distribution using BODIPY staining. Cells incubated in SLac have abundant lipid droplets distributed throughout the cell. Growing yeast in cysteine-added media promotes the mobilization of these droplets, as the staining appears distributed in the cytoplasm (Fig 10). This result is consistent with the observations made in TEM images (Supp Fig 5) and highlights potential metabolic and structural adaptations induced by cysteine in *Ustilago maydis*.

**Figure 10.**
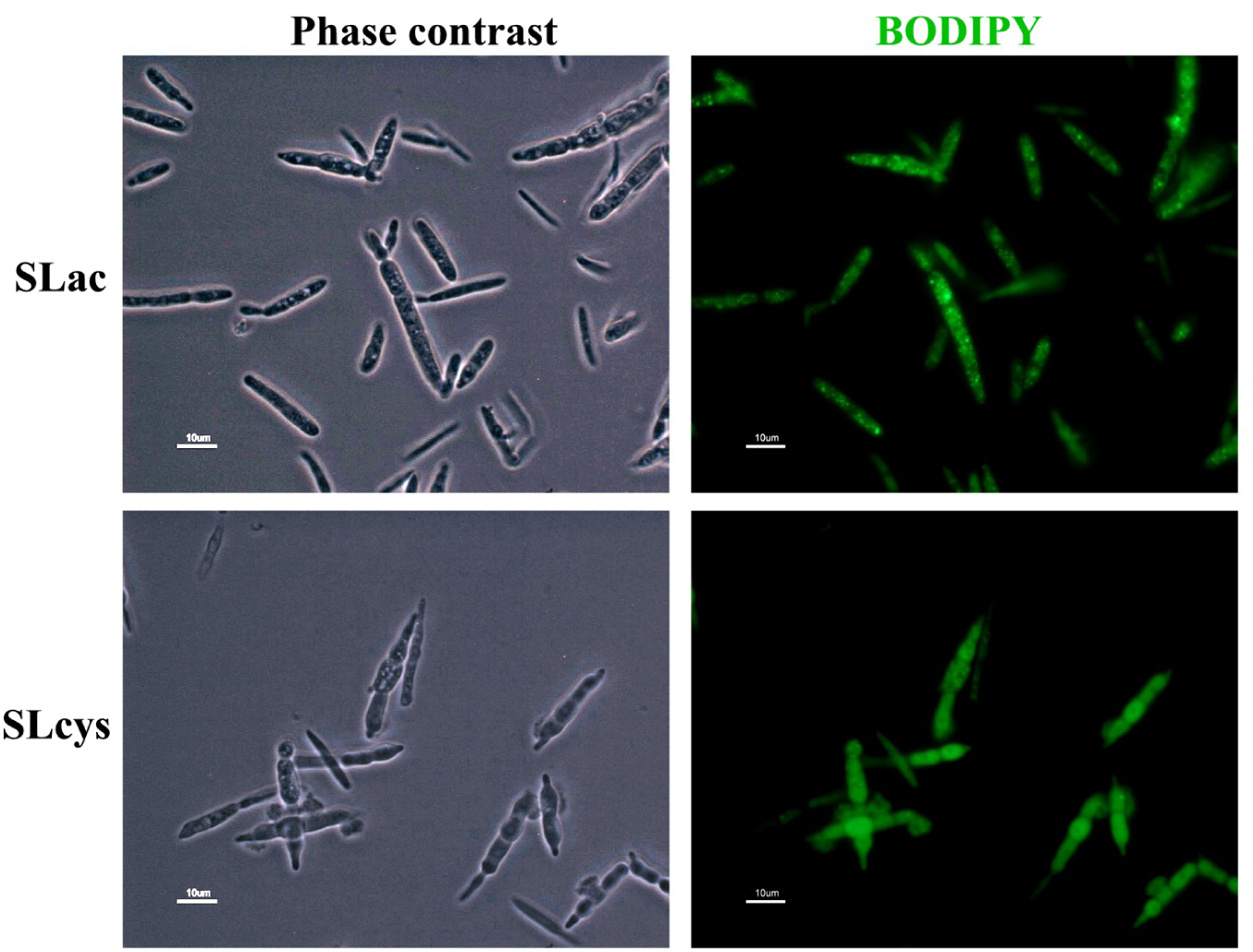
Cysteine induces lipid reorganization. *U. maydis* were stained with BODIPY and observed in an epifluorescence microscope.

### Cysteine increases protein S-sulfenylation and S-persulfidation

H_2_S induces cysteine redox modifications in several organisms[11]. In this study, we demonstrate that cysteine incubation triggers H_2_S overproduction in *Ustilago maydis*, leading to morphological and biochemical alterations in the yeast, along with an increase in its virulence. At the molecular level, H_2_S primarily drives post-translational modifications in specific cysteine residues. To further investigate these modifications, we compared cysteine PTMs between yeast grown in SLac and SLcys media using a dimedone switch [10]. Upon cysteine incubation, we observed an increase in protein S-sulfenylation (Fig. 11, right panel) and a slight increase in protein S-persulfidation (Fig. 11, middle panel). Thus, it is likely that the post-translational modifications induced by H_2_S overproduction play a regulatory role in the previously described morphological and biochemical changes.

**Figure 11.**
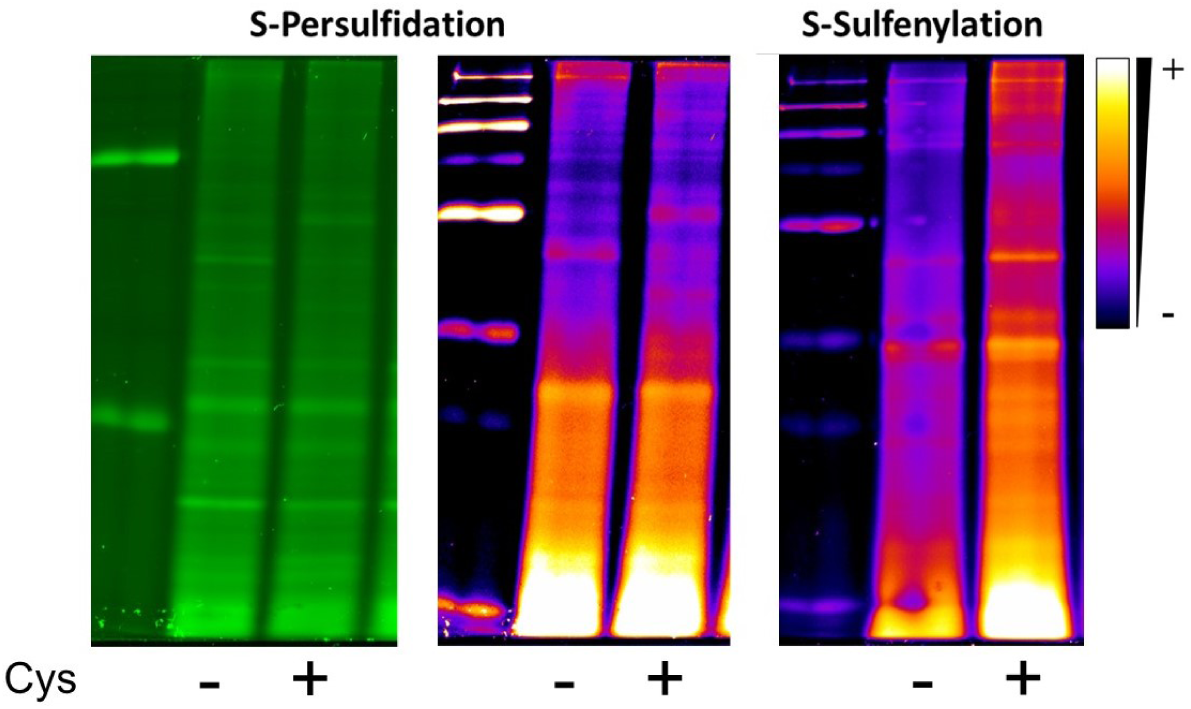
Cysteine induces PTM in *U. maydis*. Cells were cultured in SLac (-) and SLcys (+) and S-persulfidation (middle panel) and S-sulfenylation (right panel) were detected using the dimedone switch. In the left panel, total NBF-Cl stained proteins are shown in green, serving as a loading control.

## Discussion

In recent years, the relationship between H₂S metabolism and several physiological processes has been demonstrated, including photosynthesis and fruit ripening in plants [64], activation of the immune response [8], and infection mechanisms caused by bacteria [17] and fungi [32]. This gasotransmitter is produced in the cytoplasm by enzymes such as sulfite reductase, cystathionine beta-synthase (CBS), cystathionine gamma-lyase (GSE), 3-mercaptopyruvate sulfur transferase (MPST), among others. At the mitochondrial level, it is produced by different enzymes, such as sulfide quinone oxide reductase (SQOR) and cysteine desulfurase (NFS1). In *U. maydis*, several hypothetical enzymes related to sulfur metabolism have been described, such as cysteine synthase, cystathionine gamma-synthase, homoserine O-acetyltransferase, sulfite reductase, 3-mercaptopyruvate sulfurtransferase, sulfide quinone oxidoreductase, and sulfate adenylyltransferase, among others [7]. The presence of these enzymes suggests that the metabolic pathways for sulfate assimilation and transsulfuration, which include reactions leading to H₂S production, are conserved in *U. maydis*.

H₂S has been described as having a hormetic role in fungal infection. The addition of the H₂S donor sodium hydrosulfide (NaHS) at concentrations above 0.1 mM reduces fruit decay mediated by the fungi *Aspergillus niger* and *Penicillium italicum* by inhibiting spore germination, germ tube elongation, and mycelial growth [29]. A similar phenomenon has been demonstrated in the phytopathogenic fungi *Rhizopus nigricans*, *Mucor roxianus*, and *Geotrichum candidum* [65]. Conversely, inhibiting endogenous H₂S production decreases fungal infectivity, as demonstrated in *Candida albicans*, where deletion of the CBS enzyme led to delayed cell growth, reduced antioxidant response, impaired hyphal growth, and increased beta-glucan exposure [32].

In addition to its key role in fungal infection processes, H₂S-mediated protein S-persulfidation is also important in host defense mechanisms. This has been demonstrated in human hematopoietic cells, where a single nucleotide polymorphism in the gene encoding the enzyme CSE decreases H₂S production, leading to reduced levels of protein S-persulfidation and, ultimately, increased susceptibility to infection by the fungus *Aspergillus fumigatus* [31]. It was also observed that *Aspergillus* strains with low H₂S production are more susceptible to being eliminated by host cells and display reduced virulence in mice.

In plants, the enzymes L-cysteine desulfhydrase (DES1), cytosolic O-acetylserine B (OAS-B), and beta-cyanoalanine synthase (CAS-C1) produce H₂S in the cytoplasm, plastids, and mitochondria, respectively. It has been described that during the early stages of infection, plants increase DES1 activity and H₂S production to inhibit the growth of pathogens. Strategies such as fumigating plants with NaHS solutions have been implemented, particularly to inhibit the growth of fungi like *Rhizopus nigricans*, *Mucor rouxianus*, and *Geotrichum candidum* [66].

Thus, sulfur metabolism, and in particular, H₂S production, plays an important role during infection, both in plants and pathogens. This work reviewed the role of cysteine in the pathogenesis of *U. maydis* on *Z. mays* plants. The use of exogenous H₂S donors (NaHS, GYY4137, etc.) has been documented [25], nonetheless, these conditions may be an artifact and do not examine the endogenous synthesis of this gasotransmitter. In *S. cerevisiae*, it has been described that nitrogen restriction induces H₂S production by increasing the expression of genes related to the “sulfate reduction sequence” (SRS) [62]. Additionally, it has been shown that incubation in cysteine-containing media induces H₂S production [67]. Based on this background, the conditions required to stimulate H₂S production in *U. maydis* FB-2 and SG200 strains were investigated. H₂S production was evaluated in fermentative media (YPD and SD) and respiratory media (YPLac and SLac) with and without 2 mM cysteine (Fig. 1A and 1B). Under these conditions, we found that H₂S production significantly increases in respiratory media supplemented with cysteine in both strains (Supp Fig 1). Regarding H₂S production, it begins at around 6 hours of culture (Fig. 1B), leading to significant accumulation at 24 hours (Fig. 1A).

The induction of H₂S production associated with an increase in cysteine concentration is a conserved phenomenon, from yeasts [1] to mammalian cells [68]. Regarding the carbon source, it is interesting that H₂S production increases in respiratory media, as the overproduction of this gasotransmitter has only been observed in the presence of glucose in fermentative yeasts like *S. cerevisiae*. This phenomenon suggests a relationship between H₂S production and mitochondrial metabolism. In mammals, H₂S can be produced in mitochondria by enzymes such as MPST and function as an electron donor in the respiratory chain through its interaction with the protein sulfide quinone oxidoreductase (SQR). In yeast, NFS1 or cysteine desulfurase, is an essential gene whose protein Nfs1 is responsible for mitochondrial H_2_S production and is involved in iron-sulfur cluster assembly [69]. Thus, H₂S production plays a role in maintaining mitochondrial metabolism, reminiscent of the metabolic conditions when the atmosphere was reductive [70]. These results suggest a relationship between H₂S production and mitochondrial function. To evaluate the mitochondrial response to endogenous H₂S production induced by cysteine, the relative mitochondrial abundance was assessed using fluorescence microscopy and transmission electron microscopy. Incubation in respiratory culture media induces mitochondrial biogenesis, and this effect is enhanced when the media is supplemented with cysteine (Fig. 6). Transmission electron microscopy results suggest that cysteine induces mitochondrial fusion, particularly in SLcys, where elongated mitochondria were observed (Fig. 7). Thus, incubation in nitrogen-restricted respiratory media (SLcys) induces mitochondrial biogenesis and H₂S production. This result is interesting because it has been shown that H₂S regulates mitochondrial function depending on its concentration. At low concentrations, it fails to induce mitophagy but stimulates mitochondrial biogenesis and stress response. At optimal concentrations, mitochondrial homeostasis is maintained, while in excess, H₂S inhibits mitochondrial activity by blocking the cytochrome pathway of the respiratory chain [71]. Therefore, it is likely that incubation in SLcys not only alters mitochondrial morphology but also affects mitochondrial energy function, leading to changes in energy production and/or reactive oxygen species (ROS) generation. To explore this further, oxygen consumption was measured in yeast cells grown in SLac or SLcys. Basal or endogenous respiration was slightly higher in cells grown in the presence of cysteine, suggesting increased respiratory activity. When glucose was added to evaluate oxygen consumption in response to a carbon source, we observed that cells pre-incubated with cysteine further increased cellular respiration, suggesting an optimized respiratory chain. It has been described that excess H₂S inhibits the activity of complex IV of the respiratory chain [72]. To evaluate this, oxygen consumption was measured in the presence of cyanide, which inhibits complex IV activity. In cells grown without cysteine, oxygen consumption was inhibited, suggesting that the respiratory chain depends on complex IV or the cytochrome pathway. In contrast, when cells were grown in the presence of cysteine, oxygen consumption was not inhibited by cyanide (Fig. 8). Thus, the inhibition of complex IV by H₂S is inducing adaptive mechanisms to maintain mitochondrial metabolism, primarily through the activity of an alternative oxidase, which has already been described as being expressed when *U. maydis* is subjected to stress conditions [73]. To determine if cyanide-resistant respiration depends on an alternative oxidase, respiration was measured using n-Octyl Gallate, and complete inhibition was observed, indicating that when cells are grown in the presence of cysteine, complex IV is inhibited, and alternative oxidase activity increases. This is consistent with studies showing that the alternative oxidase of *U. maydis* increases the metabolic plasticity of the cells [74].

Another effect of modulating complex IV activity in the respiratory chain is the dysregulation of ROS production, a phenomenon conserved in mammals [75] and yeast [76]. To determine the effect of the described conditions on ROS production in *U. maydis*, H₂O₂ production was measured using the Amplex Red assay. SLcys cultured yeast have an increased content of H_2_O_2_ in the supernatant and also increased H_2_O_2_ production. These results suggest that H₂S production induced by growth in cysteine increases ROS production, either stimulating H₂O₂ production or decreasing the activity of cellular antioxidant systems (Fig. 9).

Protein synthesis is highly dynamic and sensitive to changes in nutrient quantity and type in yeast. In *S. cerevisiae*, it has been described that under glucose-limiting conditions, more than 50 proteins related to the use of alternative carbon sources, beta-oxidation, and oxidative phosphorylation are upregulated. In contrast, under nitrogen-limiting conditions, more than 50 proteins related to scavenging alternative nitrogen sources and protein degradation are upregulated [77]. Another study described that glucose restriction modifies the overexpression of 183 proteins related to metabolism or respiration [78].

In *U. maydis*, the stress response (NaCl, sorbitol, and heat stress) was evaluated using 2D electrophoresis and proteomics. The proteins that increased expression include cytoplasmic and mitochondrial heat shock proteins, amino acid metabolism enzymes, oxidative phosphorylation pathway enzymes, protein translation machinery, and mitochondrial metabolism enzymes [79]. The overproduction of ROS is closely related to the overproduction of H₂S. The effect of ROS on protein homeostasis is unclear, but it suggested that oxidative stress induces cytosolic aggregation of ubiquitin-related proteins in *S. cerevisiae*. This phenomenon suggests that ROS produced under stress conditions can also modulate other processes, such as protein degradation or homeostasis.

Consequently, we decided to evaluate the protein profile changes induced by cysteine using the label-free system. Around 17,609 peptides and 1706 proteins were detected, among which at least 400 proteins showed differential expression between the treatments (Fig. 4). The distribution of the identified proteins based on expression changes and p-value is shown in the volcano plot (Fig. 3). Regarding the proteins with at least a two-fold increase in expression, 190 were overexpressed and were grouped based on KEGG pathway, biological process, or local network cluster string. Among the overexpressed proteins, the most represented cluster was related to mitochondrial function (Krebs cycle enzymes, mitochondrial carriers, and respiratory complex proteins, among others). The second group of proteins with altered expression are related to protein degradation via the proteasome (26S proteasome regulatory subunit RPN2, proteasome subunit beta, and proteasome endopeptidase complex) and nitrogen recycling (aspartate aminotransferase, homoisocitrate dehydrogenase, ornithine aminotransferase, threonine dehydratase, and arginine biosynthesis bifunctional protein ArgJ among others). Therefore, a complex protein related to protein translation regulation (Eukaryotic translation initiation factor 3) was upregulated. Some reports showed that septins play a role in fungal pathogenesis [80] [81] [82]. Here, we found an increase in proteins from the septin GTPase family in yeast grown in SLcys (Fig 5).

On the other hand, several cytoplasmic proteins were underexpressed upon cysteine incubation, including metabolic enzymes such as S-adenosylmethionine synthase, adenosylhomocysteinase, pyruvate dehydrogenase, hsp90 cochaperone, glycerol-3-phosphate dehydrogenase, arginase, glutamate dehydrogenase, and lipoyl synthase, as well as antioxidant enzymes like peroxiredoxin and superoxide dismutase (Fig 5). Thus, this protein profile suggests a decrease in antioxidant response that correlates with higher H_2_O_2_ production in *U. maydis*.

*U. maydis* pathogenesis depends on several factors, like beta-oxidation enzymes. Strains lacking hydroxyacyl coenzyme A dehydrogenase showed a decreased virulence, suggesting a role of mitochondrial lipid metabolism on pathogenesis [83]. In this work, yeast grown in SLcys showed more than four times the expression of this enzyme, probably increasing *U. maydis* virulence. Other enzymes related to lipid metabolism with higher expression are Very-long-chain 3-oxoacyl-CoA reductase, Very-long-chain (3R)-3-hydroxyacyl-CoA-dehydratase, and aldehyde dehydrogenase. To determine if the described conditions affect lipid distribution, a lipid stain with BODIPY was performed. We observed that incubation in media with cysteine resulted in less accumulation of lipid droplets and more pronounced cytoplasmic staining, suggesting increased lipid mobilization (Fig. 9). These results were consistent with images observed in TEM microscopy (Supp Fig 5). Experiments conducted in *S. cerevisiae* showed that cysteine addition upregulates the expression of genes primarily associated with amino acid metabolism and, to a lesser extent, carbohydrate metabolism. The link between cysteine metabolism and lipid metabolism has been documented. In HepG2 cells, cysteine exposure reduces the expression of SREBP, FAS, G6PDH, and SCD1 genes, leading to a parallel decrease in triglyceride concentration [84]. In the hepatic tissue of rats on a high-fat diet, cysteine supplementation induces the expression of malic enzyme, fatty acid synthase, and hydroxymethylglutaryl-CoA reductase, also increasing cholesterol and triglyceride levels [85].

It has been documented that pathogenic fungi increase their infective capacity when subjected to stress conditions [86]. In *U. maydis*, it has been described that defects in genes encoding enzymes involved in beta-oxidation reduce the fungus’s infective capacity [87] [88].

Thus, we decided to evaluate whether the metabolic modifications induced by cysteine influence the infectivity of the fungus on maize plants. The results obtained indicate that 12 days post-infection, plants infected with *U. maydis* pre-incubated in SLac showed a 31.6% tumor formation rate, while crops infected with SLcys developed tumors in 46.3% of the plants. In plants, anthocyanins synthesis serves as a response mechanism to oxidative stress [89]. In this study, we observed an increase in anthocyanins synthesis when maize plants were infected with yeast grown in SLcys (Fig 2 and Supplementary Figure 3).

### Conclusions

The results obtained in this article suggest that cysteine metabolism and probably H₂S production play a role in the pathogenicity of *U. maydis*. A likely mechanism involves the overproduction of H₂S, which, by inhibiting the activity of complex IV in the respiratory chain, stimulates the generation of H₂O₂ and other stress mechanisms that eventually lead to increased fungal infectivity. *Ustilago maydis* is a phytopathogenic fungus that induces losses in maize crops; however, in recent years, the controlled cultivation of this fungus for gastronomic purposes has been promoted [90].

In recent years, interest has arisen in developing strategies to control fungal growth without involving the use of antifungals, particularly due to the increase in resistance to these drugs [91] [92]. Thus, there is a need to search for metabolism regulators that modify the infective capacity of organisms. It is likely that among sulfur metabolism intermediates, inhibitors of H₂S production or fumigation of plants with synthetic or natural H₂S donors [93] could be an alternative in agriculture.

### Supporting information

**S1 Fig.** Cell viability upon incubation in SLac or SLcys. Cells were grown under the above-mentioned conditions for 24 hrs, then spotted onto YPD agar plates and incubated for an additional 48 hrs.

**S2 Fig**. H₂S production is similar between the strains. *Ustilago maydis* strains SG200 and FB2 were inoculated in SLac and SLcys media, and H₂S production was assessed using the lead acetate strip method after 24 hours.

**S3 Fig.** Cysteine increases *U. maydis* pathogenicity. Yeast cells cultivated under the above-mentioned conditions were inoculated on maize plants and an increase in tumor formation, chlorosis, and anthocyanins production was observed.

**S4 Fig.** Catalase activity is reduced in yeast grown in cysteine. Twenty-four hours after the aforementioned culture conditions, H_2_O_2_ elimination was measured (p<0.001).

**S5 Fig.** Cysteine induces lipid redistribution in *U. maydis*. Yeast incubated as mentioned above were analyzed in TEM. Lipid droplets appear as white spherical structures within the cells. Panel A (SLac) showed more lipid droplets than panel B (SLcys). The figure is magnified 8000 x.

**S1 Table.** Upregulated proteins in SLCys treated yeast.

**S2 Table.** Downregulated proteins in SLCys treated yeast.

## Acknowledgments

This work was funded by DGAPA-PAPIIT-UNAM México, Grant numbers: FTQ: IN208922 and IN220125; NTR: IA204723 and IN214625 and by CONAHCYT Mexico FTQ: CF-2023-I-884, LVT and MMJ: IPN-SIP: 20240946, 20242205. DRR is a CONAHCYT fellow CVU 1083449 enrolled in the Ciencias Bioquímicas PhD program at UNAM. Mass spectrometry experiments were conducted during the first theoretical-practical course on mass spectrometry “Proteómica Basada en Espectrometría de Masas”, organized by CINVESTAV Zacatenco and the Mexican Proteomics Society (SMP). Special thanks are extended to Bc. Sc. Lorena Ramírez Reyes and PhD. Josaphat Miguel Montero Vargas for their technical support. We also acknowledge the technical support provided by PhD Lilia Morales-Garcia, Bc Sc Alma Olivia Sánchez González and PhD. Jesús Aguirre Linares in the oximetry experiments. We are grateful for the technical support provided by PhD. Teresa Lara Ortíz and PhD. Gabriel del Río Guerra in the use of the TECAN equipment. Also, we would like to thank to PhD. Laura Ongay Larios, BSc Maria Guadalupe Codiz Huerta, and MSc Minerva Mora Cabrera from the IFC-Molecular Biology Unit for their technical support. Finally, the authors want to thank Manuel Ortinez and Aurey Galvan from the maintenance workshop, and Juan Barbosa and Ivett Rosas from the Computing Unit for their continuous assistance.

## Author contributions

**E.E.-S.** conducted the H₂S production experiments, mass spectrometry (MS), fluorescence microscopy, biochemical measurements, and oximetry experiments. **N.T.-R., R.O.-H., and O.E.-M.** performed fluorescence and electron microscopy. **M.J.-M. and L.V.-TK** conducted the infection assays. **E.R.-C.** assisted with and performed the MS experiments. **D.R.-R.** carried out the S-persulfide and S-sulfenyl detection assays. **F.T.-Q.** designed the research, analyzed the data, and wrote the paper.

---

**Supplementary figure 1.**
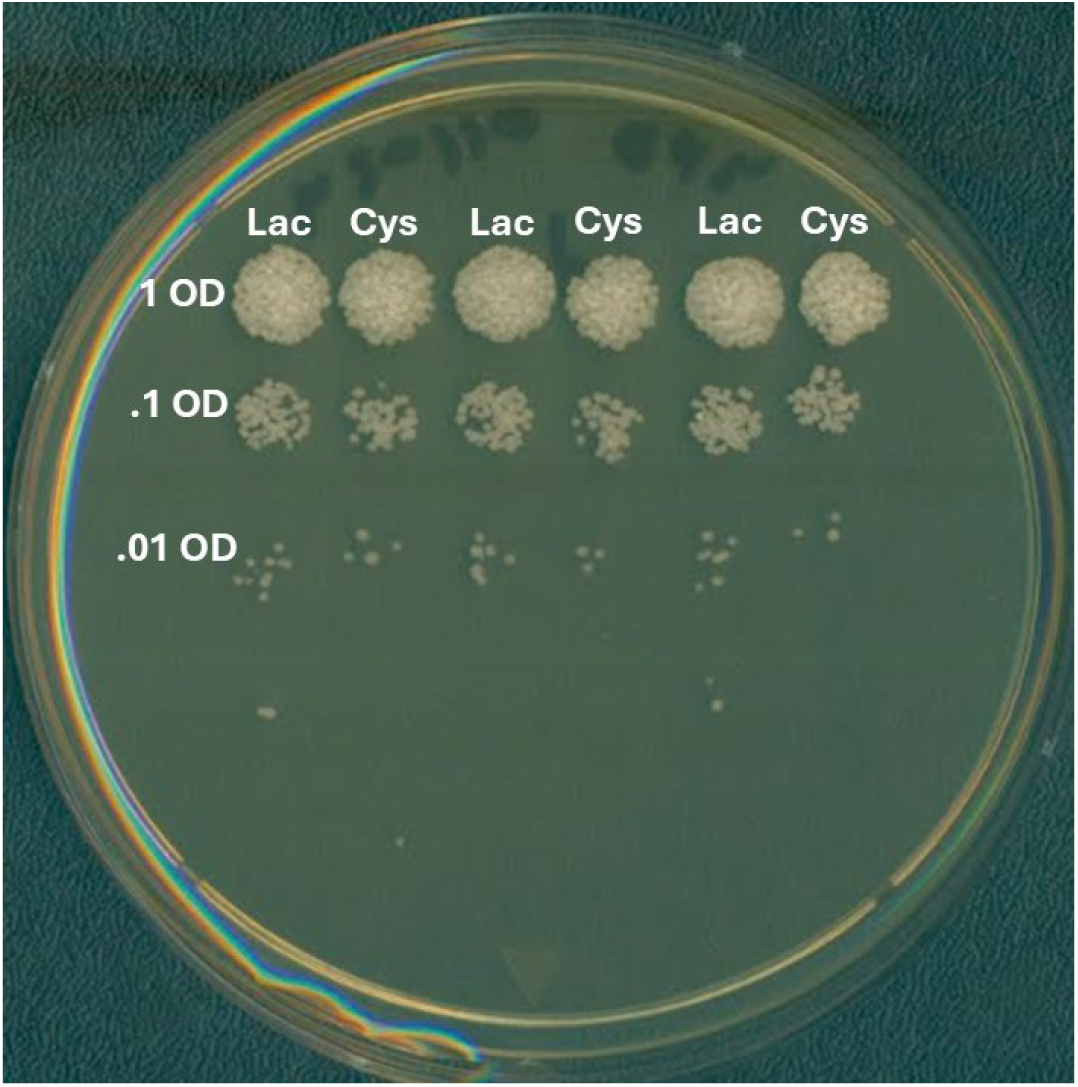
Cell viability upon incubation in SLac or SLcys. Cells were grown under the above-mentioned conditions for 24 hrs, then spotted onto YPD agar plates and incubated for an additional 48 hrs.

**Supplementary figure 2.**
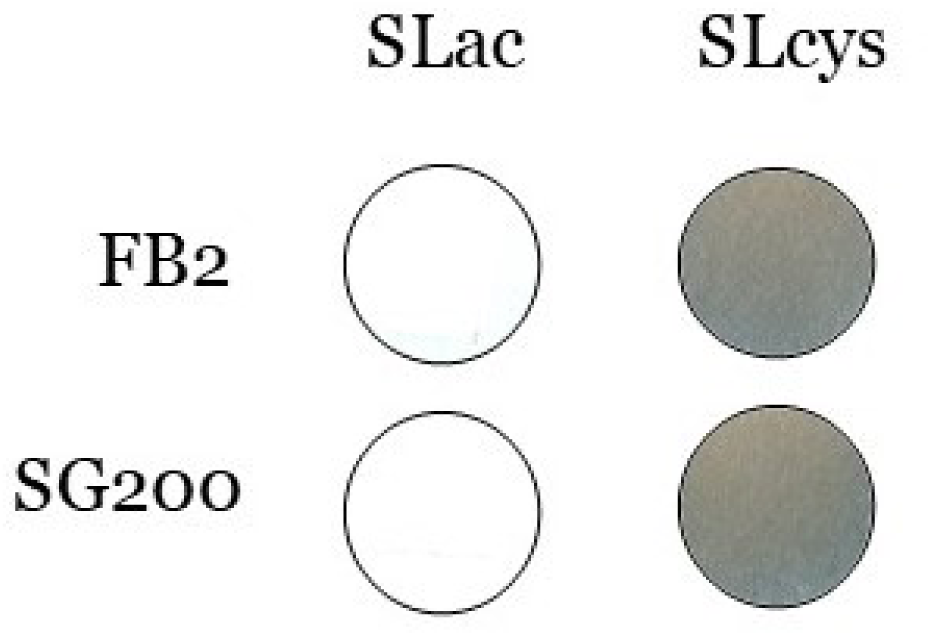
H₂S production is similar between the strains. *Ustilago maydis* strains SG200 and FB2 were inoculated in SLac and SLcys media, and H₂S production was assessed using the lead acetate strip method after 24 hours.

**Supplementary figure 3.**
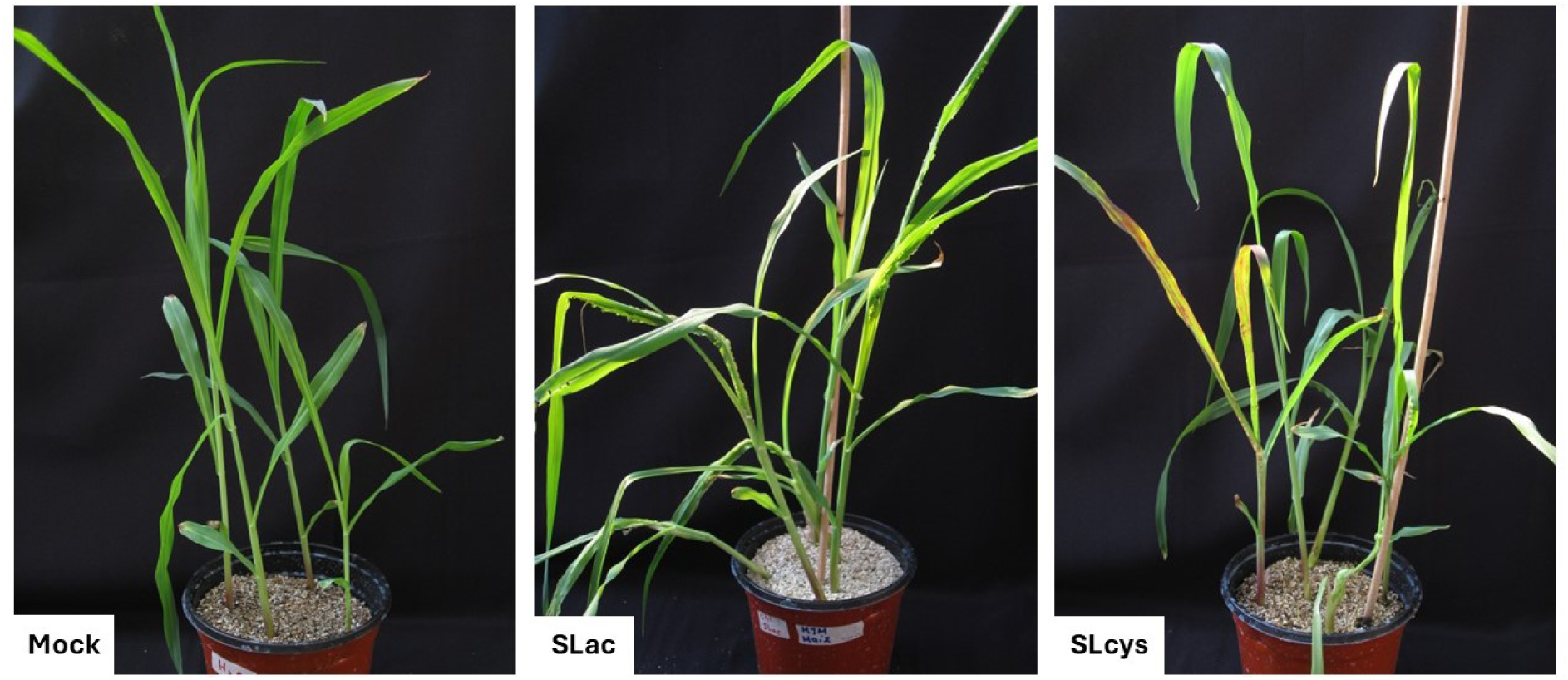
Cysteine increases *U. maydis* pathogenicity. Yeast cells cultivated under the above-mentioned conditions were inoculated on maize plants and an increase in tumor formation, chlorosis, and anthocyanins production was observed.

**Supplementary figure 4.**
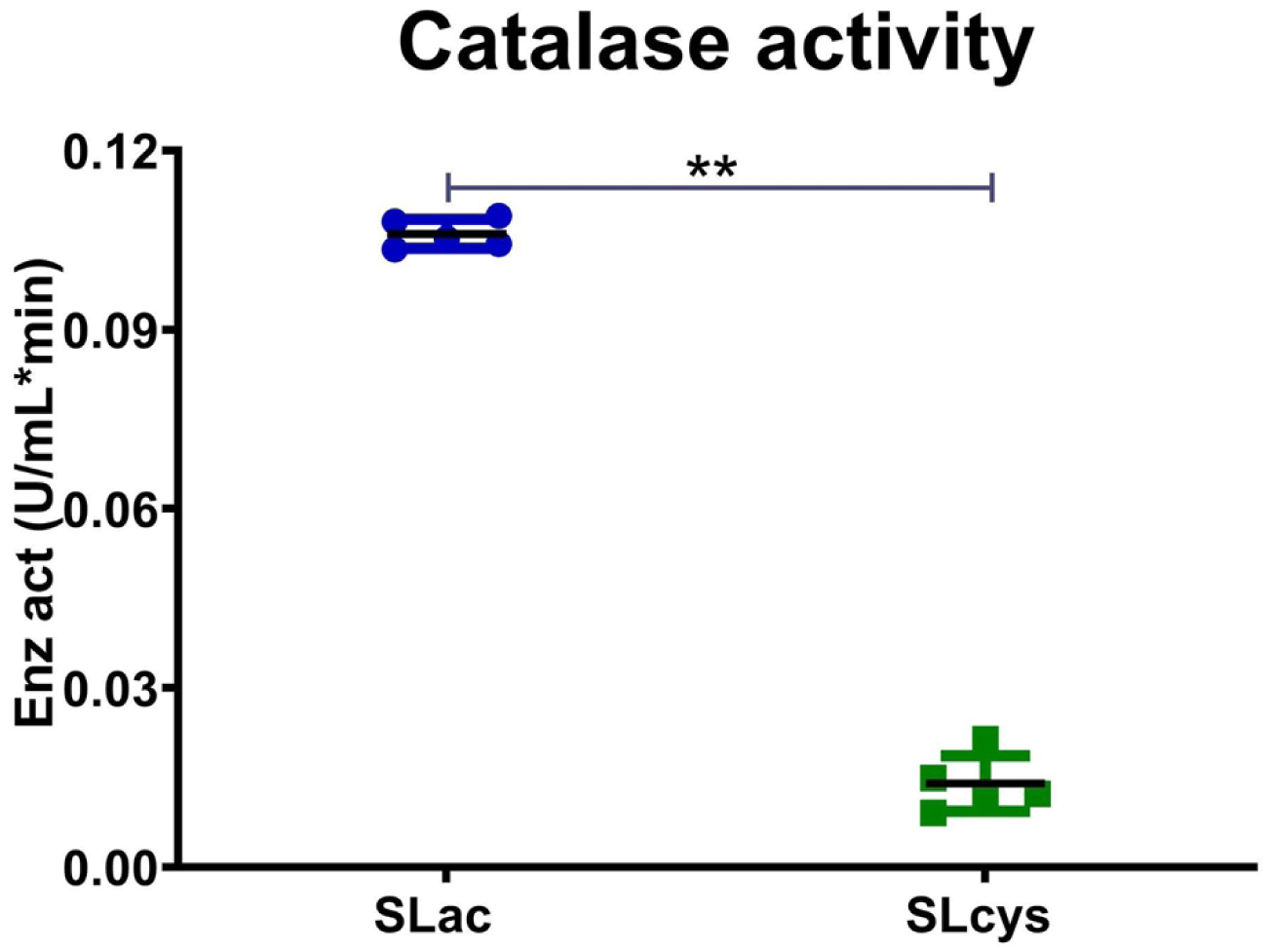
Catalase activity is reduced in yeast grown in cysteine. Twenty-four hours after the aforementioned culture conditions, H_2_O_2_ elimination was measured (p<0.001).

**Supplementary figure 5.**
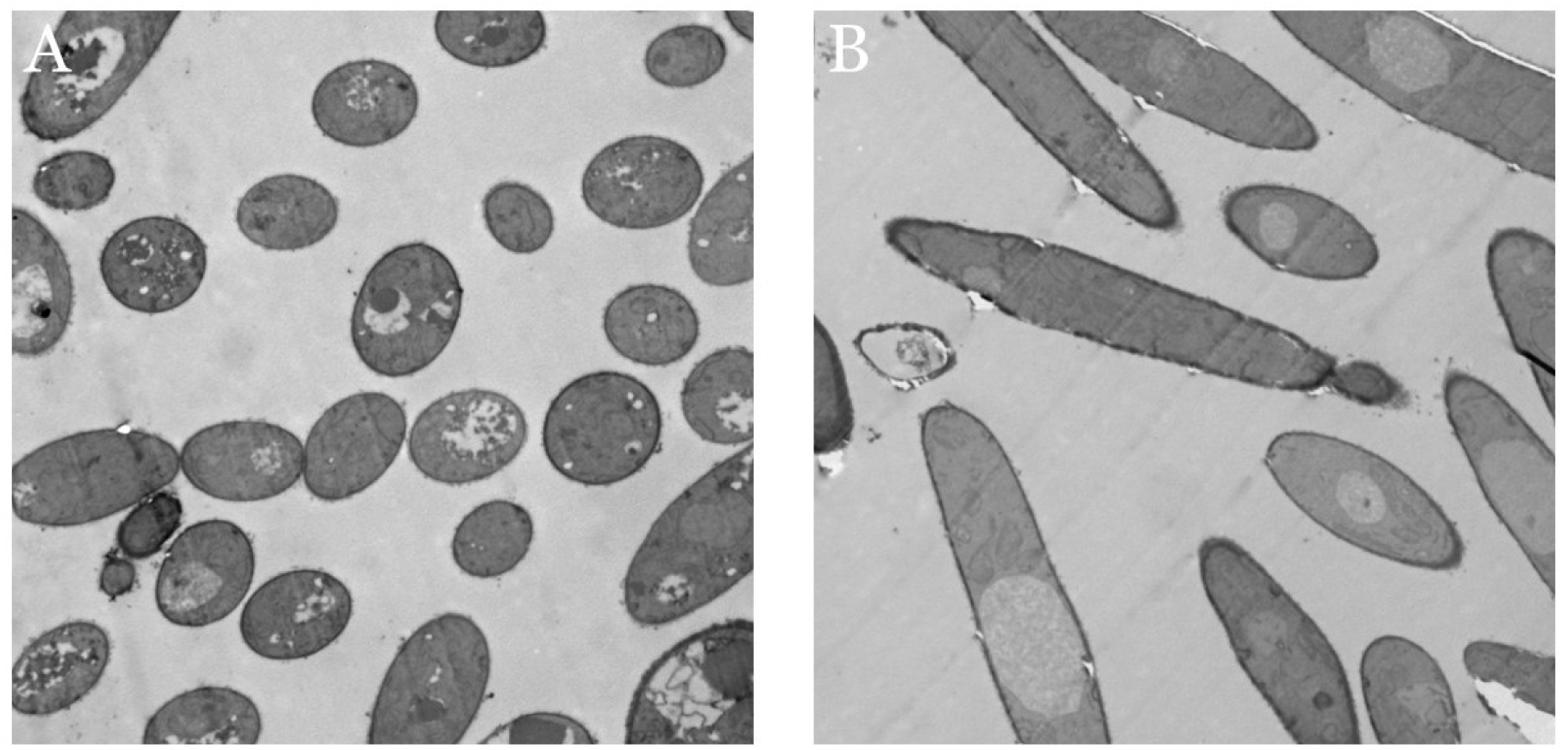
Cysteine induces lipid redistribution in *U. maydis*. Yeast incubated as mentioned above were analyzed in TEM. Lipid droplets appear as white spherical structures within the cells. Panel A (SLac) showed more lipid droplets than panel B (SLcys). The figure is magnified 8000 x.

Supplementary table 1. Upregulated proteins in SLCys treated yeast: Supplementary table 1 <u>March25.xlsx</u>

Supplementary table 2. Downregulated proteins in SLCys treated yeast Supplementary table 2 <u>March25.xlsx</u>

